# Vitamin B5 Metabolism is Essential for Vacuolar and Mitochondrial Integrity and Xenobiotic Detoxification in Fungi

**DOI:** 10.1101/2023.12.10.571002

**Authors:** Jae-Yeon Choi, Shalev Gihaz, Muhammad Munshi, Pallavi Singh, Pratap Vydyam, Patrice Hamel, Emily M. Adams, Kevin Fuller, Choukri Ben Mamoun

## Abstract

Fungal infections, a leading cause of mortality among eukaryotic pathogens, pose a growing global health threat due to the rise of drug-resistant strains. New therapeutic strategies are urgently needed to combat this challenge. The PCA pathway for biosynthesis of Co-enzyme A (CoA) and Acetyl-CoA (AcCoA) from vitamin B5 (pantothenic acid) has been validated as an excellent target for the development of new antimicrobials against fungi and protozoa. The pathway regulates key cellular processes including metabolism of fatty acids, amino acids, sterols, and heme. In this study, we provide genetic evidence that disruption of the PCA pathway results in a significant alteration in the susceptibility of fungi to a wide range of xenobiotics, including clinically approved antifungal drugs through alteration of vacuolar integrity and drug detoxification. The drug potentiation-mediated by genetic regulation of genes in the PCA pathway could be recapitulated using the pantazine analog PZ-2891 as well as the celecoxib derivative, AR-12 through inhibition of fungal AcCoA synthase activity. Collectively, the data validate the PCA pathway as a suitable target for enhancing the efficacy and safety of current antifungal therapies.

## INTRODUCTION

Annually, invasive fungal infections kill over 1.6 million people globally. A 2023 report by the US Center for Disease Control (CDC) warned that the rapid growth of incidence of infection by multi-drug resistant *Candida auris* urgently threatens domestic and global public health ^1^. Azole-resistant *Aspergillus* spp. ^2,3^ (classified by the CDC as an emerging threat) and other drug-resistant strains ^4^ contribute to the urgency for development of new antifungal therapeutic strategies.

Development of highly selective and potent antifungal drugs (AFDs) is hampered by the similarities shared between mammalian and fungal cells and the latter’s drug resistance mechanisms ^5–7^. One of these commonalities lies in the central importance of the synthesis of Coenzyme-A (CoA), an obligate cofactor of approximately 4-9% of all known enzymes ^8^ and a precursor for acetyl-CoA (AcCoA), an acetyl carrier essential for the operation of synthetic and oxidative pathways. The biosynthesis of CoA involves a 5-step enzymatic process that begins with the phosphorylation of vitamin B5 (pantothenate or PA) by pantothenate kinases (PanKs) to form 4-phosphopantothenate. Various components of the PA-CoA-AcCoA (PCA) pathway (**Fig. 1A**) have previously been explored as potential druggable targets for antimicrobials, including against malaria parasites ^9,10^. Human cells express four pantothenate kinases (PANK1α, PANK1β, PANK2, and PANK3) in different tissues. Mutations in the *PANK2* gene, encoding a mitochondrial PanK enzyme, have been linked to a debilitating pediatric neurodegenerative disease named pantothenate-kinase associated neurodegeneration (PKAN) ^11^. Modulation of the cytoplasmic PANK3 activity using small molecule modulators to compensate for the loss of PANK2 has become of interest in the treatment of PKAN. In contrast, in *Saccharomyces cerevisiae* as well as most causative agents of invasive fungal infections, a single copy of the PanK enzyme performs the first step in the PCA pathway ^12,13^. Fungal *PANK* genes have all been shown to be essential for cell viability as complete or conditional knockouts of these genes lead to cell death ^12–16^. As a result, efforts to identify inhibitors of fungal PanK enzymes has been considered an attractive strategy for the development of new antifungal drugs. A high throughput screen of a library of 156,593 chemical compounds against the *Aspergillus fumigatus* PanK enzyme (AfPanK) identified several pyrimidone triazoles as potential strong and selective inhibitors of AfPanK activity ^16^. The first crystal structure of a fungal PanK enzyme was obtained at 1.8 Å resolution for the *S. cerevisiae* PanK enzyme, Cab1p, encoded by the *CAB1* gene ^16^. It consists of two monomers sharing a dimeric interface that accommodates the ATP-binding pocket, active site, and part of the PA binding domain. The structure of the enzyme was also determined in the presence of pyrimidone triazoles inhibitors and used for molecular docking with other known modulators of PanK including the pantazine PZ-2891, an orthosteric modulator of human PanK3 ^16,17^. An analog of PZ-2891, BBP-671, is being studied in a human clinical trial (NCT04836494) for the treatment of PKAN.

**Fig. 1.**
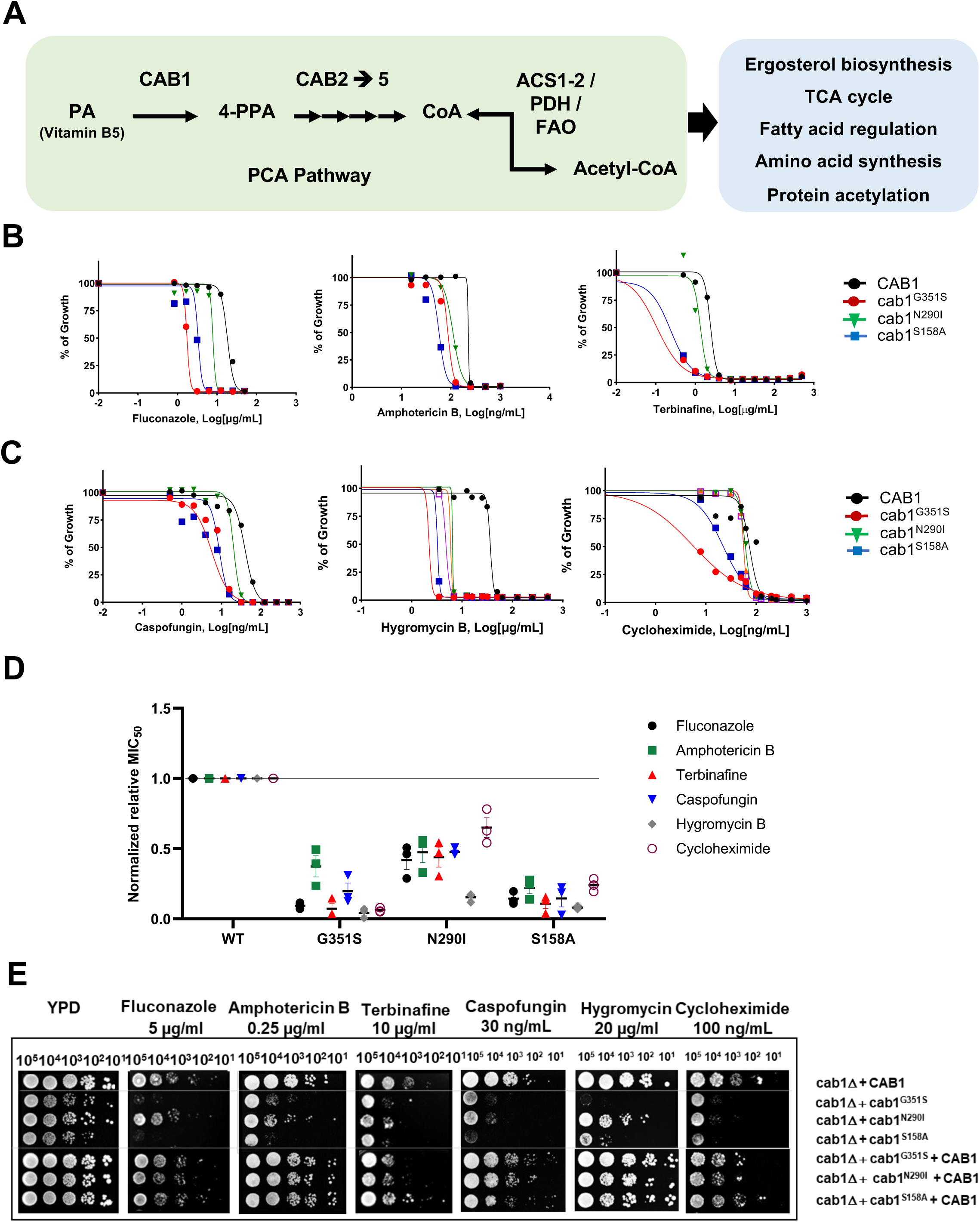
*Cab1* mutants display growth defects and increased susceptibility toward ergosterol targeting drugs. **(A)** Schematic diagram of Coenzyme A synthetic pathway and biological roles of the Acetyl-CoA. Pantothenate kinase (PanK) catalyzes the phosphorylation of pantothenic acid to form 4’-phosphopantothenic acid, the first step in the biosynthesis of CoA. Phosphopantothenoylcysteine synthase (PPCS) converts 4’- phosphopantothenic acid to phosphopantothenoylcysteine. Phosphopantothenoylcysteine decarboxylase (PPCD) catalyzes the decarboxylation of phosphopantothenoylcysteine to form pantetheine. Pantetheine kinase (PANK) catalyzes the phosphorylation of pantetheine to form dephospho-CoA. Dephospho-CoA kinase (DPCK) catalyzes the transfer of a phosphate group from ATP to dephospho-CoA, resulting in the formation of CoA. **(B and C)** The *cab1* mutants display susceptibility toward drugs targeting ergosterol pathway and non-ergosterol targeting pathway. Yeast liquid growth curve assay with known AFDs targeting the ergosterol biosynthesis pathway was done using yeast cells harboring different *CAB1* mutants. Cells were inoculated into 100 µL of YPD liquid media containing the antifungals in serial dilutions at 10 cells per µL ratio and incubated at 30°C while cell growth was monitored by OD_600_. % of cell growth of individual mutant strain in presence of AFDs was obtained compared to the cell growth in the vehicle control. (**D)** Normalized relative MIC_50_ of yeast strains producing *Cab1* variants against AFDs. MIC_50_ values were determined on MIC_50_ data obtained from the liquid growth assays presented in Figure 1B and 1C. These values were calculated by dividing the MIC_50_ value of the *cab1* mutant or the wild type strain for the specified AFD by the MIC_50_ value of the wild type strain. (**E**) Complementation of the wild type *CAB1* gene restores AFD resistance in the *cab1* mutants to the similar level as that observed in wild-type cells. The *cab1Δ/w303-1B* strain harboring 1) wild type *CAB1*, 2) *cab1^mutant^*, or 3) *cab1^mutant^* + wild type *CAB1*, were inoculated into synthetic glucose medium and grown overnight. Cells were harvested and re-suspended in 0.9% NaCl. Ten-fold serial dilutions of cells were spotted onto the YPD plates containing fluconazole (5 µg/mL), amphotericin B (0.25 µg/mL), and terbinafine (10 µg/mL), caspofungin (30 ng/mL), hygromycin (20 µg/mL), or cycloheximide (100 ng/mL), and incubated at 30°C for 3 days.

The first evidence linking PanK activity to drug susceptibility was reported by Chiu et al. using a *S. cerevisiae cab1^ts^* thermosensitive mutant carrying a mutated chromosomal *CAB1* gene. The *cab1^ts^* thermosensitive strain (JS91.14-24) was generated following EMS mutagenesis to select mutants altered in saturated fatty acid biosynthesis ^13,18^. The *cab1^ts^* mutant harbors, among multiple other genetic mutations (40 identified following whole genome sequence (unpublished data), a mutated Cab1 enzyme with 4 individual substitutions, one of which, G351S, is directly linked to the thermosensitive phenotype ^13^. The mutant was found to be susceptible to amphotericin B and tolerant to fluconazole, amorolfine and terbinafine ^12^. Recent studies have linked the susceptibility to amphotericin B directly to the G351S mutation, whereas tolerance to fluconazole, amorolfine and terbinafine was caused by other mutations in other genes unrelated to the PCA pathway (unpublished data). To gain further insights into the link between the PCA pathway and antifungal drug susceptibility and to unravel the underlying molecular mechanisms, we used strains that carry a chromosomal deletion of *CAB1* but carry either the wild type or mutated alleles of *CAB1* on a centromeric plasmid ^14–16^. Our analyses revealed that specific mutations in *CAB1* cause susceptibility to a broad-spectrum of antifungals, including to those that target ergosterol biosynthesis and others acting on varied cellular processes (protein synthesis, cell wall formation, and RNA synthesis). Cell biological and pharmacological analyses established a role for CoA biosynthesis from pantothenate in the regulation of vacuolar function, including the detoxification of xenobiotics. We identified the PanK3 orthosteric activator, PZ-2891, and the celecoxib derivative, AR-12, as modulators of the PCA pathway and enhancers of fungal susceptibility to antifungal drugs.

## RESULTS

### PanK mutants have broad-range susceptibility to antifungal drugs

Previous studies in *S. cerevisiae* and *A. fumigatus* have shown that reduced PanK activity caused by substitution of glycine 351 to serine (cab1^G351S^) in the *S. cerevisiae* Cab1 enzyme, or reduced expression, using a conditional promoter to drive the expression of the *A. fumigatus* AfPanK, result in altered susceptibility to amphotericin B and voriconazole, respectively ^12,16^. To gain further insights into the link between Cab1 activity, the PCA pathway (**Fig. 1A**), and fungal susceptibility to antifungal drugs, a detailed characterization of *S. cerevisiae* drug susceptibility using strains carrying various mutations in the *CAB1* gene was conducted. The yeast strains used in these studies carry a chromosomal deletion of the *CAB1* gene but express on a centromeric plasmid either a wild type *CAB1* (*cab1Δ+CAB1*), mutant alleles of *CAB1* (*cab1Δ+cab1^m^* with m= *cab1^G351S^*, *cab1^N290I^*, *and cab1^S158A^*) or both the mutant alleles and the wild type *CAB1* (add-back: *cab1Δ+cab1^m^+CAB1*). As shown **Fig. 1B**, all the yeast strains expressing different *CAB1* alleles exhibited increased susceptibility to fluconazole, amphotericin B, and terbinafine, compared to the isogenic strain carrying the wild type *CAB1*. The *cab1Δ+cab1^G351S^* mutant was found to be highly susceptible to all ergosterol biosynthesis inhibitors examined with a reduction in MIC_50_ values determined to be ∼12, ∼15, and ∼3-fold for fluconazole, terbinafine, and amphotericin, respectively, compared to the wild type strains (**Fig. 1D**). Similarly, the *cab1Δ+cab1^S158A^* mutant displayed reduced MIC_50_ values with fold reductions compared to the wild type determined to be ∼7, ∼9, and ∼5 for fluconazole, terbinafine, and amphotericin, respectively (**Fig. 1D**).

The broad-range susceptibility of *cab1* mutants to antifungal drugs led us to examine whether the underlying mechanism could be linked to disruption of a particular metabolic process such as ergosterol biosynthesis by the PCA pathway or to a broader disruption of yeast’s ability to detoxify xenobiotics. Therefore, we examined the susceptibility of the mutants to drugs that target unrelated pathways including hygromycin and cycloheximide, which target protein synthesis, and caspofungin, which targets cell wall integrity (**Fig. 1C**). Similar to their susceptibility to ergosterol biosynthesis inhibitors, the *cab1* mutant alleles showed higher susceptibility to caspofungin, hygromycin and cycloheximide compared to the wild type (**Fig. 1C**). The *cab1^G351S^* mutation resulted in the highest drug susceptibility with the MIC_50_ values of caspofungin, hygromycin, and cycloheximide determined to be ∼5, ∼23, and ∼16-fold lower compared to the wild type (**Fig. 1D**). Similarly, *cab1^S158A^* mutation resulted in reduced MIC_50_ values for caspofungin, hygromycin, and cycloheximide by ∼6, ∼13, and ∼4-fold compared to the wild type (**Fig. 1D**). All complemented strains carrying the wild-type *CAB1* gene displayed susceptibility levels comparable to those of the wild-type (WT) strain (Fig. 1E). Altogether these data demonstrate that inhibition of PanK activity leads to enhanced susceptibility to a wide variety of antifungal drugs.

### PanK-deficient cells have altered vacuole biogenesis and xenobiotic detoxification mechanism

The overall enhanced drug susceptibility of Cab1 defective mutants led us to investigate a possible role of the vacuole in this mechanism. In fungi, the vacuole plays a critical role in the detoxification of xenobiotics such as drugs and metals ^19^ ^12^. Therefore, we examined whether the *cab1* mutants might also be susceptible to metals such as FeSO_4_ and CuSO_4_, which are detoxified in the vacuole. Consistent with an altered vacuolar detoxification in the mutants, the growth of the *cab1Δ+cab1^G351S^*, *cab1Δ+cab1^N290I^*and *cab1Δ+cab1^S158A^* mutants was dramatically reduced in media supplemented with FeSO_4_ or CuSO_4_ compared to the wild type and complemented strains (**Fig. 2A**). Vacuolar integrity was further investigated by measuring the accumulation of the cell-tracker dye, CMAC, using fluorescence microscopy ^19,20^. The *cab1Δ+cab1^G351S^*, *cab1Δ+cab1^N290I^* and *cab1Δ+cab1^S158A^* mutants were all found to have enlarged vacuoles compared to the wild type and complemented strains (**Fig. 2B and Fig. 2C**), with the vacuoles in the mutants occupying 60 to 70% more of the total cell area compared to the vacuoles of the wild-type and complemented strains. Vacuolar enlargement in the mutants was further confirmed by electron microscopy as shown in **Fig. 2D**. Taken together, these data provide the first evidence that pantothenate phosphorylation regulates vacuolar homeostasis and xenobiotic detoxification.

**Fig. 2.**
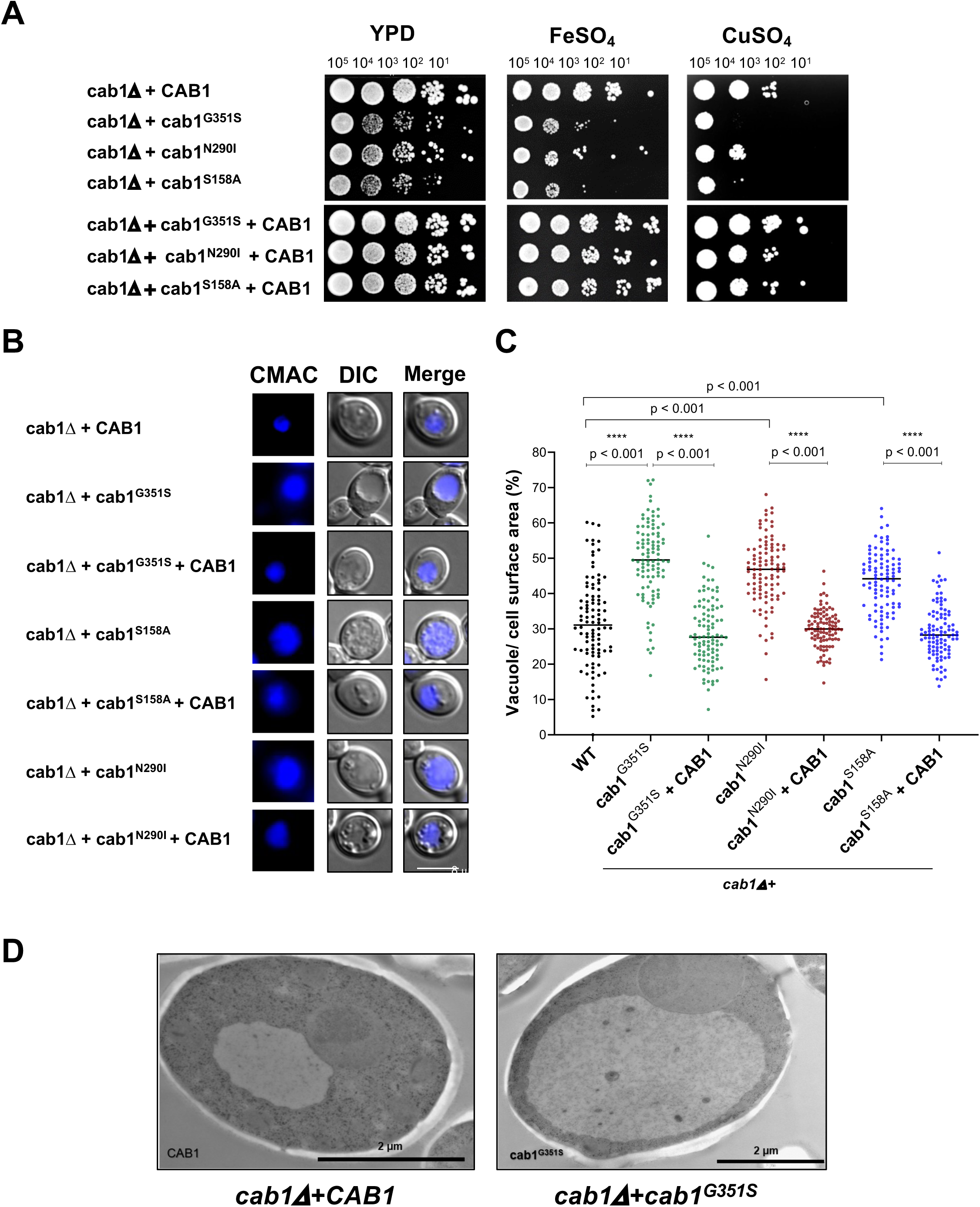
Yeast strains expressing various *CAB1* mutations have defects in detoxification and vacuolar function and structure. **(A)** The *cab1* mutations altered yeast capacity to overcome metal toxicity. Solid growth assays were performed with the yeast strains described above in glucose media supplemented with 7 mM FeSO_4_ or with 10 mM CuSO_4_. **(B)** Morphological analysis of vacuolar defects shows that *cab1* mutant strains have unusually enlarged vacuoles. Immuno-fluorescence microscopy images of *cab1Δ* + *cab1*^mutant^ strains described above show enlarged and/or fragmented vacuoles, while the *cab1Δ* + *CAB1*^WT^ and the *cab1Δ* + *cab1* ^mutant^ + *CAB1*^WT^ strains have typical vacuolar morphology. **(C)** Quantitative analysis of vacuolar area size (as a proportion of total cell area) as a function of *CAB1* status. **(D)** Electron microscopy images of *cab1Δ* + *CAB1^WT^* and *cab1Δ*+*cab1*^G351S^ single cells confirm enlargement of vacuole induced by G351S mutation.

Recent studies have established an important role for vacuolar biogenesis in the maintenance of mitochondrial function and integrity ^21,22^. Therefore, we surmised that altered vacuolar function in the *cab1* mutants could also result in altered mitochondrial function. Accordingly, the *cab1Δ+cab1^G351S^*, *cab1Δ+cab1^N290I^*and *cab1Δ+cab1^S158A^* mutants showed severe growth defects on non-fermentable carbon sources (glycerol, ethanol, and lactate-based media) (**Fig. 3A**) and altered oxygen consumption rates (OCR) (**Fig. 3B-E**). Consistent with these findings, immunofluorescence assays aimed to localize the mitochondrial outer membrane protein Por1p revealed that unlike the wild-type and complemented strains, the *cab1* mutants exhibited fragmented mitochondria (**Fig. 3F**). Since the overproduction of reactive oxygen species (ROS) is associated with dysfunctional mitochondria ^23,24^, cellular ROS levels of the *cab1* mutants were also determined by measuring the conversion of the non-fluorescent dihydrorhodamine 123 (DHR-123) to the fluorescent rhodamine 123. As shown in **Fig. 3G**, ROS levels in the *cab1Δ+cab1^G351S^*, *cab1Δ+cab1^N290I^* and *cab1Δ+cab1^S158A^* mutants were found to be 10.1, 13.0 and 4.1-fold higher than in the wild-type and complemented strains, respectively.

**Fig. 3.**
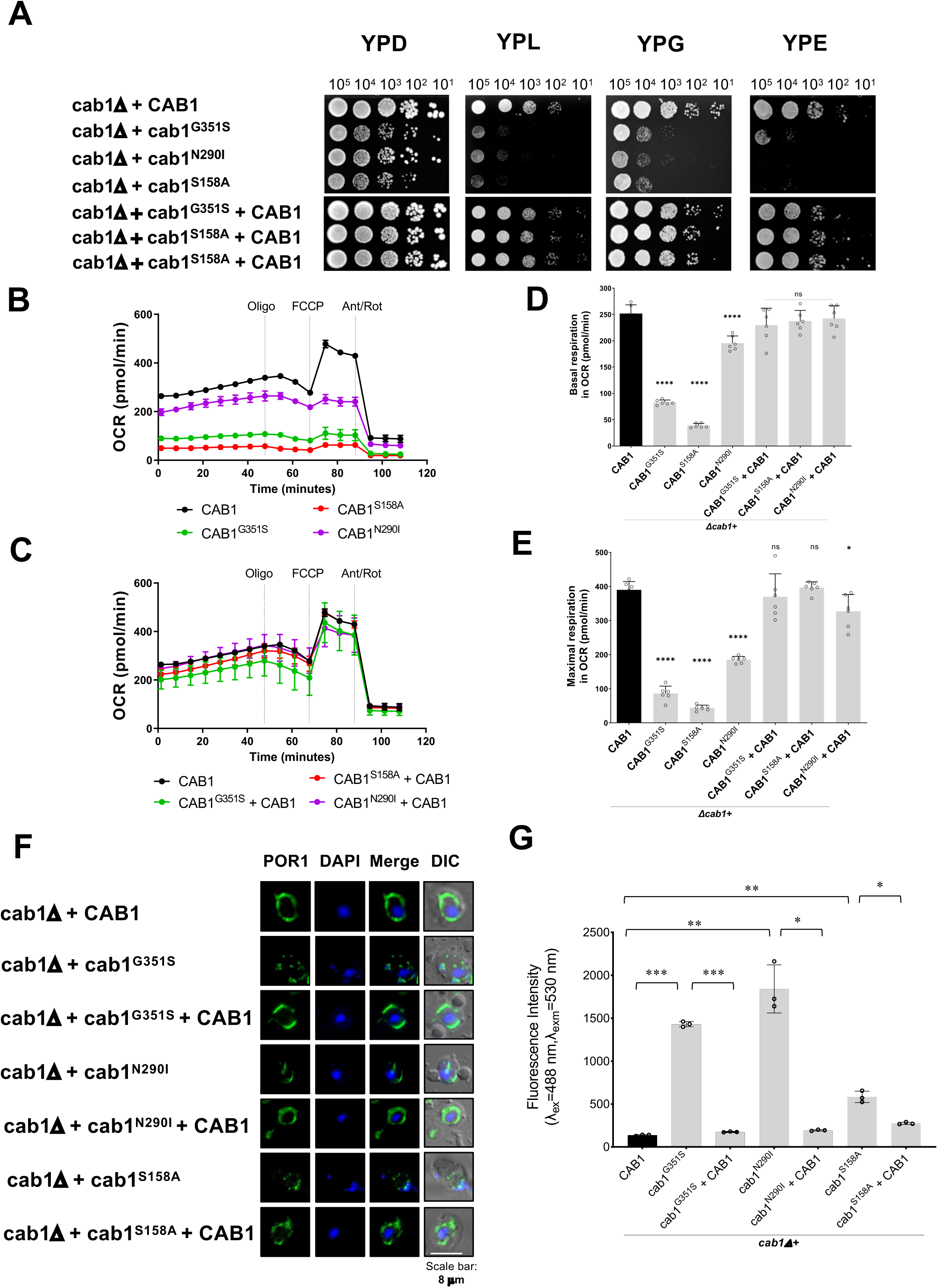
Yeast *cab1Δ* strains harboring various *CAB1* mutations display defects in mitochondrial function. **(A)** Solid growth assays reveal that mutations in *CAB1* alter strains’ ability to utilize non-fermentable carbon sources. Solid growth assays using ten-fold serial dilutions plated onto YPD (glucose), YPL (lactate), YPG (glycerol), or YPE (ethanol) media and observed over 3-4 days. **(B)-(C)** Oxygen consumption rate (OCR) of yeast cells harboring *CAB1* variants. OCR profile of *cab1Δ* strains using Seahorse 96X and Mito Stress kit. Dashed lines represent the injections of mitochondrial uncoupling drugs; oligomycin, FCCP, antimycin A, and rotenone. **(D)-(E)** Basal respiration and maximal respiration of *cab1Δ* strains (t-tests performed for each group comparing mutants to the wild type strain at *p=0.05*). **(F)** *cab1Δ* strains exhibit mitochondrial structural defects. Immuno-fluorescence microscopy images of *cab1Δ* strains reveal aberrant mitochondria structures. **(G)** *cab1Δ* strains have increased levels of reactive oxygen species (ROS). ROS analysis was performed using dihydrorodamine 123.

### Altered pantothenate phosphorylation leads to reduced pantothenate utilization and CoA biosynthesis and increased cysteine levels in yeast

To gain further insights into the mechanism by which altered pantothenate phosphorylation leads to defective vacuolar biogenesis, we assessed the effect of the Cab1 mutations on the activity of the PCA pathway. Endogenous pantothenate utilization of *cab1Δ*+*cab1^G351S^*, *cab1Δ*+*cab1^S158A^*, and *cab1Δ*+*cab1^N290I^* was determined by measuring following metabolism of ^14^C-pantothenate (Fig. 4A). As shown in Fig. 4A, all *cab1* mutants showed significantly lower levels of ^14^C-pantothenate utilization compared to the isogenic strain carrying the wild type *CAB1*. The lowest pantothenate utilization level (∼2% that of the wild type) was measured for the *cab1^S158A^* mutant followed by 25% for the *cab1^G351S^*mutant and 34% for the *cab1^N290I^* mutant. Expression of the wild-type *CAB1* gene in these strains restored PA utilization to levels similar or above those in the isogenic wild type-strain. Consistent with the reduced pantothenate utilization in the mutants, cellular CoA levels in the mutants were also significantly lower compared to the wild-type and complemented strains (Fig. 4B and Fig. S1). Because reduced pantothenate phosphorylation results in less phosphopantothenate available for the second step in CoA biosynthesis catalyzed by phosphopantothenoylcysteine synthetase (Cab2) to form phosphopantothenoylcysteine from phosphopantothenate and cysteine (Fig. 4D), we reasoned that altered Cab1 activity would also result in accumulation of cysteine. As shown in Fig. 4C, cysteine levels increased by 2.7, 3.3, and 2.2-fold in the *cab1^G351S^*, *cab1^S158A^*, and *cab1^N290I^* mutants, respectively, compared to the wild-type or complemented strains. Consistent with these findings, analysis of the transcription profile of the *cab1^G351S^*, *cab1^S158A^*, and *cab1^N290I^* mutants showed a significant downregulation of the genes involved in sulfur assimilation and the cysteine/methionine biosynthetic pathway compared to the wild type and mutant strains carrying a wild type *CAB1* gene (Fig. 4D-E and Fig. S2 and S3). Among these genes, the expression of the ATP sulfurylase-encoding gene, *MET3,* α-subunit of sulfite reductase, Met10, cystathionine β-synthase, Cys4, bifunctional dehydrogenase and ferrochelatase, MET8, and sulfate permease, SUL2, genes in each of the mutants decreased dramatically (between 75% and ∼95%) compared to the wild type strain (Fig. 4E and S2).

**Fig. 4.**
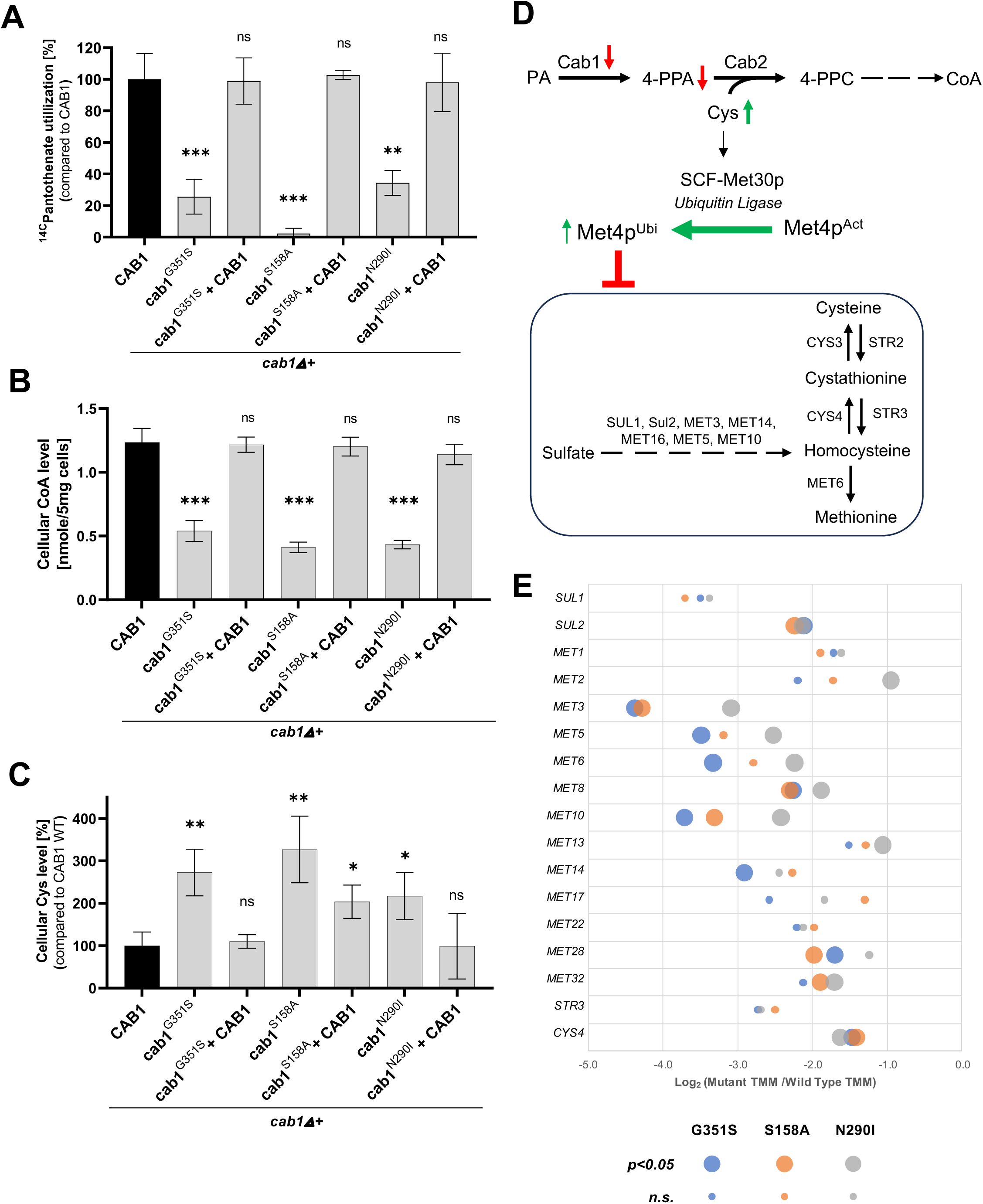
Metabolic defects in yeast strains harboring *cab1* mutations. **(A)** PA utilization in the *cab1Δ* strains harboring various *CAB1* mutations. Cell free extracts of yeast *cab1* mutants were used to measure the endogenous PA utilization of *cab1Δ + CAB1*, *cab1Δ + cab1^G351S^*, *cab1Δ + cab1^G351S^*+ *CAB1*, *cab1Δ + cab1^S158A^*, *cab1Δ + cab1^S158A^ + CAB1*, and *cab1Δ + cab1^N290I^*, *cab1Δ + cab1^N290I^ + CAB1* strains using D-[1-^14^C] pantothenate as a substrate for 10 min at 30°C. **(B)** Cellular levels of CoA in *cab1Δ* strains. CoA levels were measured using metabolite extracts from the yeast strains mentioned above grown in the presence of 0.2 µM PA. **(C)** Cysteine cellular levels of *cab1Δ* strains harboring various *CAB1* mutations. Cellular cysteine levels were measured using the metabolite extracts from the yeast strain mentioned above grown in the presence of 0.2 µM PA. For these assays, t-test was done per each group compared to the parent *CAB1* wild-type strain (*p=0.05*). **(D)** Schematic of the connection between the PCA pathway and the SUL/MET pathways. Transcription of the *SUL/MET* genes, responsible for synthesizing the crucial sulfur-containing amino acids methionine and cysteine, is mediated by the transcription factor Met4. This regulatory process is sensitive to changes in cellular cysteine levels. When cysteine levels increase, Met4 undergoes ubiquitination by the ubiquitin ligase Met30, leading to Met4’s inactivation and subsequent repression of the *SUL/MET* genes. **(E)** RNA-Seq analysis for cysteine and sulfur homeostasis expressed in *cab1Δ* strains. Expression values are TMM normalized and compared using a double-sided t-test. Large circles in corresponds to genes for which the p-value (WT vs. mutant) <0.05, but the p-value (WT vs. complement) >0.05, indicating the null hypothesis of equal expression was rejected for the mutant, but not the addback, at p=0.05. Small circles correspond to genes for which either of these criteria were not met. The gene list with annotations shown in Table S1.

### Inhibition of fungal ACS2 or V-type ATPase enzymes increases susceptibility to antifungals

The genetic data described above demonstrated a direct role of pantothenate utilization and CoA biosynthesis in yeast susceptibility to antifungals through alteration of vacuolar detoxification. Considering the potential implication of these findings on fungal therapy and reversal of multidrug resistance, we assessed whether a pharmacological approach using compounds that inhibit Cab1 activity or downstream steps such as AcCoA synthesis or vacuolar V-ATPase could mimic these genetic findings and usher in a new antifungal treatment modality. Analysis of the transcription profile of the *cab1* mutants showed a dramatic decrease (12.6%, 12.9%, and 44.6% of that of the wild type) in the expression of the *ACS1* gene encoding one of two yeast AcCoA synthetases (**Fig. S3**). Interestingly, a yeast mutant carrying the *ACS2* gene under the regulatory control of the tet-off promoter (*acs2*-tet-off) was highly susceptible to caspofungin, fluconazole and terbinafine following addition of doxycycline (**Fig. 5B**). The MIC_50_ for caspofungin shifted from 16 ng/ml in the absence of doxycycline to 10 ng/ml in the presence of the compound; that for fluconazole from 4.8 µg/ml to 0.06 µg/ml; and that for terbinafine from 4.5 µg/ml to 0.005 µg/ml. Consistent with these data, the celecoxib derivative AR-12, which is also a potent inhibitor of fungal AcCoA synthetases ^25,26^ increased yeast susceptibility to caspofungin (MIC_50_ shift from 16 ng/ml to 3 ng/ml in the absence vs presence of AR-12), fluconazole (MIC_50_ shift from 14.6 µg/ml to 0.9 µg/ml), and terbinafine (MIC_50_ shift from 3.6 µg/ml to 0.07 µg/ml) (**Fig. 5C and Fig. S4**). Finally, because of the major alteration in vacuolar function and integrity in mutants altered in Cab1 activity, we examined the susceptibility of wild-type *S. cerevisiae* to caspofungin, fluconazole, and terbinafine in the absence or presence of concanamycin A, a known inhibitor of the vacuole V-Type ATPase ^27,28^. As shown in **Fig. 5D**, treatment of yeast cells with concanamycin-A potentiates their susceptibility to caspofungin (MIC_50_ shift from 36 µg/ml to 6.5 ng/ml), fluconazole (MIC_50_ shift from 8.1 µg/ml to 5.7 µg/ml), and terbinafine (MIC_50_ shift from 3.9 µg/ml to 0.9 µg/ml).

**Fig. 5.**
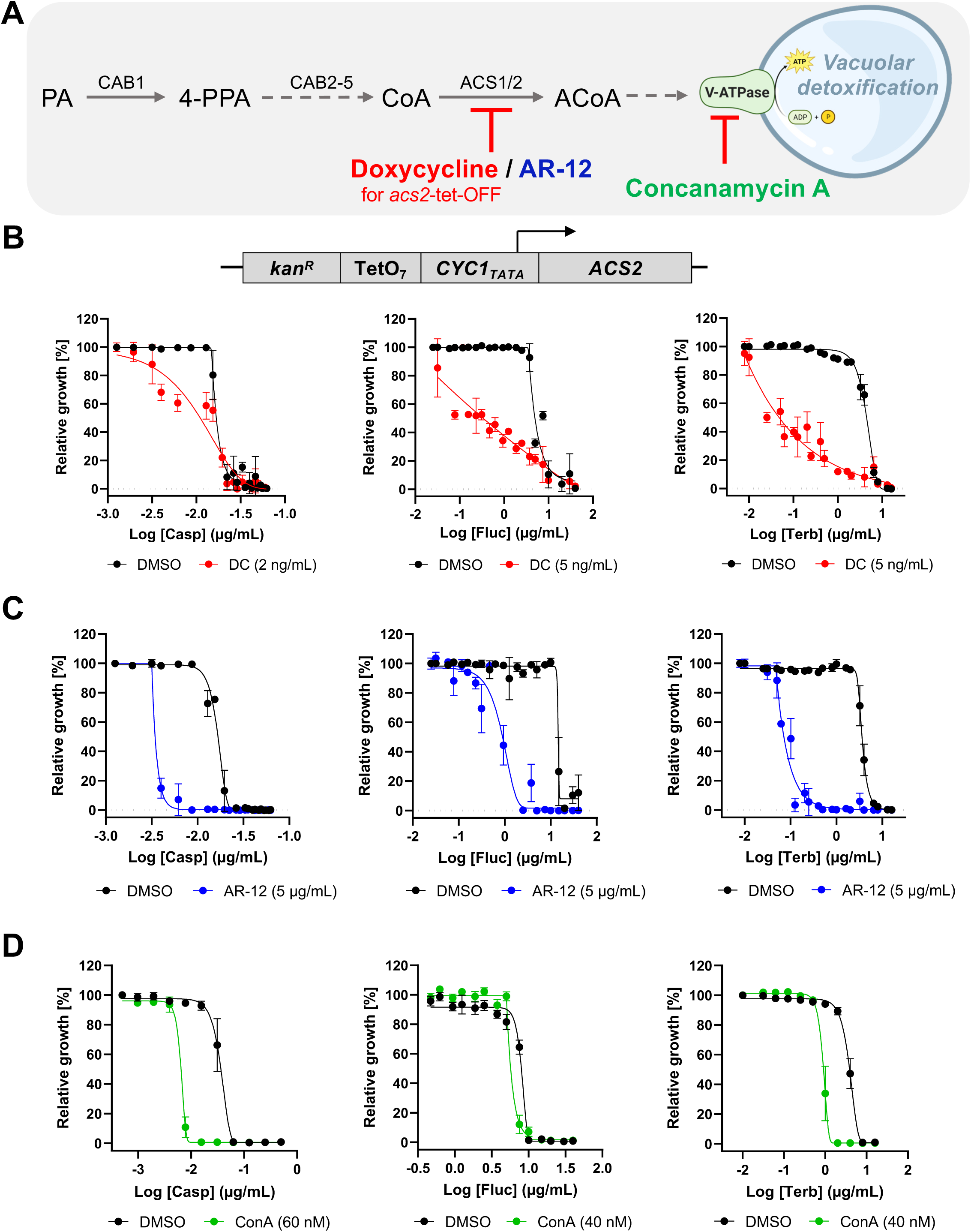
Chemical inhibition of Acs2 and V-type ATPase increases susceptibility to a variety of AFDs. **A)** Schematic of the relevant portions of the PCA pathway. **(B)** The growth of acs2 tet-off strain is inhibited by increasing concentrations of doxycycline. *S. cerevisiae* acs2 tet-off strain was inoculated in the presence or absence of doxycycline and normalized to DMSO treated wells (no drug=100% growth) and 200 µM amorolfine (0% growth). **C)** AR-12 potentiates caspofungin, fluconazole, and terbinafine by factors of ∼100x against *S. cerevisiae* WT. **(D)** Concanamycin A increases yeast susceptibility to antifungal drugs. *S. cerevisiae* WT was inoculated in the presence or absence of concanamycin A at 30°C for 48h. The growth was normalized to DMSO treated wells (no drug=100% growth) and 200 µM amorolfine well (0% growth).

### Small molecule modulation of the PCA pathway leads to increased susceptibility of pathogenic fungi to antifungal drugs

The genetic and pharmacological data described above suggest that inhibition of specific steps in the CoA biosynthesis pathway or downstream steps leading to the regulation of vacuolar detoxification could be a promising therapeutic strategy for the treatment of fungal infections to enhance the potency of approved drugs while reducing their toxicity. Therefore, we screened a library of known PanK and CoA biosynthesis modulators to search for compounds that could render pathogenic fungi susceptible to clinically approved antifungal drugs. The pantazine analog, PZ-2891, an orthosteric activator of human PANK3 ^17^, showed the highest potentiation among all compounds tested. Unlike other known Cab1 inhibitors ^12,16^, PZ-2891 did not inhibit pantothenate utilization of both WT and *cab1* mutant enzymes (**Fig. S5**). Furthermore, the compound had no antifungal activity against *S. cerevisiae*, *C. albicans* or *A. fumigatus* at concentrations up to 50 µM (**Fig. 6, Fig. S6 and Fig. S7D**). Interestingly, combinations of PZ-2891 with either amphotericin B, caspofungin or terbinafine at sublethal concentrations resulted in dramatic increases in the susceptibility of *S. cerevisiae* and *C. albicans* to these drugs (**Fig. 6A-B and Fig. S6A-B**). Similarly, PZ-2891 was found to increase the susceptibility of *A. fumigatus* to caspofungin (**Fig. 6C-D and Fig. S6C**). Unlike its inhibitory activity of human PanK3 in the absence or low levels of AcCoA, our data showed that PZ-2891 had little to no effect on Cab1 activity *in vitro* in the absence or presence of AcCoA (**Fig. S7**). Instead, the steady state levels of CoA following treatment with the compound increased by ∼1.7-fold (**Fig. 6E**). These findings suggest that the mechanism of drug potentiation mediated by PZ-2891 in yeast could be through inhibition of CoA utilization, potentially by blocking the conversion of CoA to AcCoA by Acs1, which is not essential for yeast viability on glucose-based media. Therefore, we examined the direct inhibition of Acs1 activity by PZ-2891 using a hydroxylamine-coupled assay ^26,29^. As shown in Fig. 6F, Acs1 activity was inhibited by PZ-2891 with 32.7% inhibition of the enzyme activity at 18.8 µM (**Fig. 6F**). As a control, and consistent with previous studies ^26^, the activity of purified yeast Acs1 *in vitro* was inhibited by AR-12 with a calculated IC_50_ of ∼18 µM (**Fig. 6F**). Together these data demonstrate that the mechanism of drug potentiation of the pantazine analog PZ-2891 is through alteration of a critical downstream step in the PCA pathway catalyzed by Acs1.

**Fig. 6.**
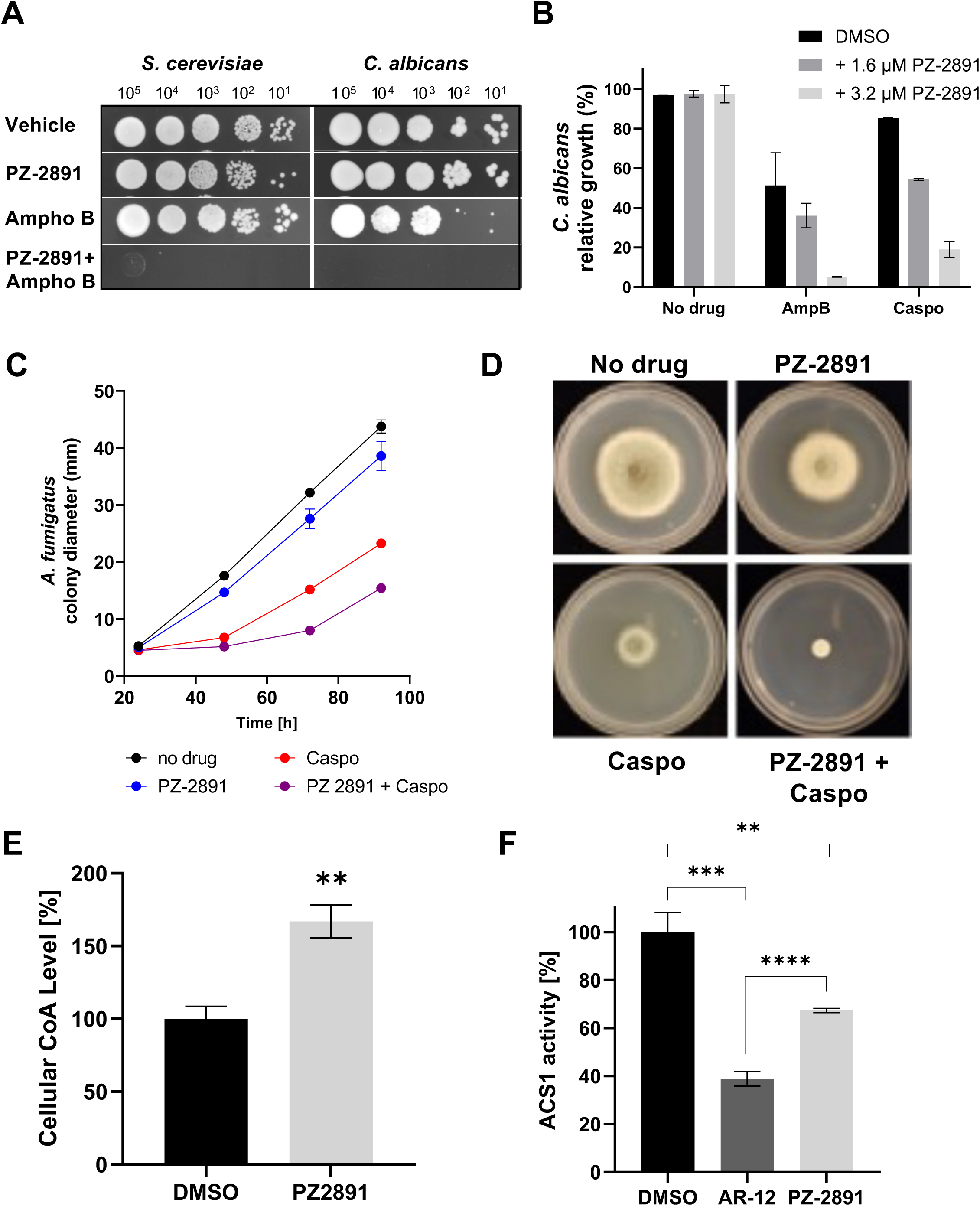
PZ-2891 increases AFD susceptibility in *S. cerevisiae*, *C. albicans* and *A. fumigatus*. **(A)** Solid growth assays showing increased *S. cerevisiae* and *C. albicans* susceptibility to amphotericin B when potentiated by PZ-2891. **(B)** Liquid growth assays showing increased *C. albicans* susceptibility to amphotericin B (125 ng/mL) and caspofungin (3.9 ng/mL) is potentiated by PZ-2891. The growth was normalized to DMSO treated wells (no drug=100% growth) and 200 µM amorolfine well (0% growth). **(C-D)** Average colony diameter rate and captures of solid media assay (up to 72h of growth) with *A. fumigatus* cells under caspofungin treatment (20 μg/ml) in the presence or absence of PZ-2891 (50 μM). **(E)** Cellular CoA levels in *S cerevisiae* following treatment with PZ-2891. CoA levels were measured using the metabolite extracts from the *S cerevisiae* cells grown in minimal glucose medium supplemented with 1 µM PA in the presence or absence of 50 µM PZ-2891. **(F)** Yeast acetyl CoA synthetase activity. The *in vitro* activity of purified enzyme from *S cerevisiae* was measured using a standard hydroxylamine-coupled assay, at 37 °C for 30 minutes, in the presence of absence of 18 µM of AR-12 or PZ-2891.

## DISCUSSION

In this study, we demonstrate that the biosynthesis of CoA from pantothenic acid and the subsequent conversion of CoA to AcCoA (the PCA pathway) play a crucial role in the regulation of vacuolar homeostasis and xenobiotic detoxification. Consequently, inhibition of the PCA pathway confers increased susceptibility to antifungal drugs, thus revealing a novel therapeutic strategy for potentiation of frontline antifungal drugs to prevent fungal infections.

Examining the implications of altered PCA pathway is fundamentally important to the understanding of the biology of fungal pathogens, and this study unveiled new cell biological mechanisms regulated by this pathway. In *S. cerevisiae*, genetic modulation of pantothenate phosphorylation unraveled major alterations in vacuolar and mitochondrial biogenesis and enhanced susceptibility to xenobiotics including metals and commonly used antifungal drugs including both drugs that target ergosterol biosynthesis inhibitors (terbinafine, fluconazole and Amphotericin B) and unrelated pathways (caspofungin, hygromycin, cycloheximide). Such broad-spectrum drug susceptibility can only be possible if major mechanisms used by fungi for drug detoxification, such as those mediated by the vacuole, are altered when the PCA pathway is inhibited. Ergosterol biosynthesis is only one of many metabolic pathways that rely on the production of AcCoA in fungi as CoA and AcCoA homeostasis also affect the TCA cycle, fatty acid regulation, amino acid synthesis, and protein acetylation ^8^. Previous reports have linked disruption in ergosterol biosynthesis to altered vacuolar ATP-powered H^+^ pumps (V-ATPase) and vacuolar acidification ^30^. It is suggested that ergosterol directly modulate the activity of V-ATPase, although the exact molecular mechanism is not yet fully understood ^30^. Our data also showed that yeast mutants with altered PanK activity can present with enlarged vacuoles, consistent with previous reports, which documented association between vacuolar dysfunction and various changes in morphology, including enlarged and fragmented vacuoles ^19,20,31–33^. Thus, it is possible that the PCA pathway controls fungal mechanisms of detoxification both through direct effect on vacuolar biogenesis and indirectly through inhibition of ergosterol biosynthesis. We propose that restricted CoA and consequently reduced AcCoA levels resulting from disruptions to the PCA pathway set off a cascade of events, leading to a deficiency in ergosterol and impaired V-ATPase function. This ultimately results in vacuolar dysfunction and subsequent loss of its ability to detoxify xenobiotics (Fig. 7).

**Fig. 7.**
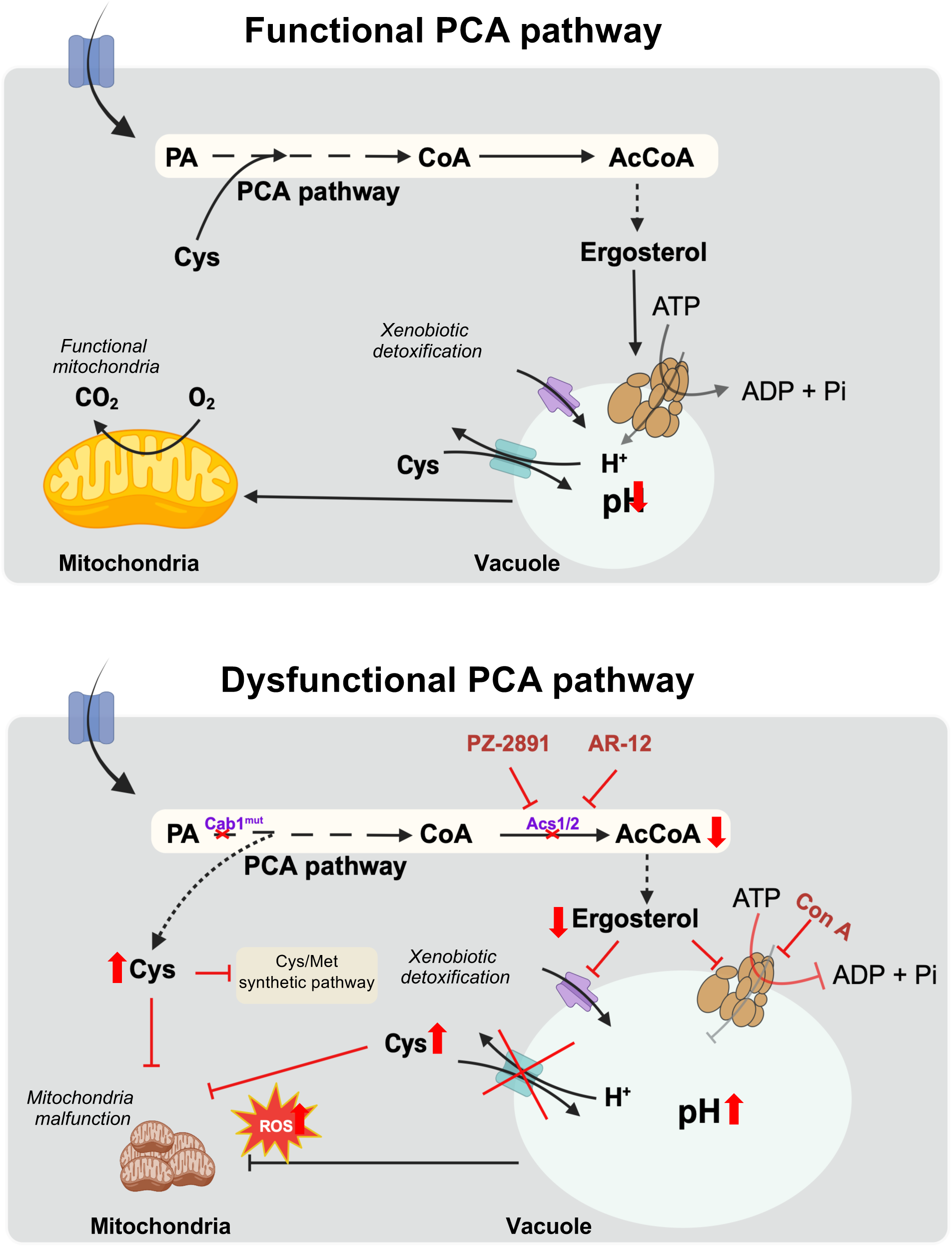
Model for PCA pathway-mediated regulation of vacuolar detoxification and susceptibility to antifungal drugs in fungi. We propose that the pathway for biosynthesis of CoA and AcCOA (the PCA pathway) from pantothenate has major regulatory function in the control of organellar biogenesis in fungi. The upper panel illustrates the cellular events associated with the functioning PCA pathway. AcCoA and ergosterol are produced through the normal PCA pathway, with ergosterol supporting the function of the V-type ATPase, thereby maintaining normal vacuole and mitochondrial function. The lower panel illustrates the dysfunctional PCA pathway, resulting in severe downstream effects on vacuolar function. Vacuolar homeostasis and xenobiotic detoxification heavily rely on the functionality of the PCA pathway in fungal strains. A reduction in ergosterol synthesis inhibits the function of the V-type ATPase, leading to an elevation of vacuolar pH, defects in cysteine sequestration, and impairments in drug/heavy metal detoxification. These vacuolar function defects also contribute to increased ROS levels and mitochondrial abnormalities. The pivotal role of the PCA pathway in these downstream cellular events creates a unique vulnerability in yeast strains that can be targeted through the use of potentiators (PAMS) like PZ-2891, in addition to other modulators of PanK, ACS, and V-ATPase enzyme activities. PA: pantothenic acid; CoA: Co-enzyme A; ROS: reactive oxygen species; PAMS: potentiators of antimicrobial susceptibility; ConA: concanamycin A.

We showed that the susceptibility of yeast cells altered in CoA biosynthesis from pantothenic acid to a broad spectrum of antifungals can be recapitulated through genetic and pharmacological inhibition of specific enzymes downstream of the PCA pathway. In yeast, AcCoA can be formed from CoA through multiple routes including by the AcCoA synthetases, Acs1 and Acs2 (See **Fig. 1A**). Cells lacking both *ACS1* and *ACS2* genes are inviable, as are cells lacking *ACS2* in glucose medium since *ACS1* is subject to glucose repression ^34,35^. Our studies demonstrated that repression of the *ACS2* gene results in increased susceptibility to commonly used antifungal drugs. This phenotype was further replicated using the Acs1/2 inhibitor AR-12 and the V-type ATPase inhibitor concanamycin A.

Unlike its function as an activator of the human pantothenate kinases (PANK3), we found that the pantazine PZ-2891 has major drug potentiation activity in fungal cells. Growth assays demonstrated potentiation of both caspofungin and amphotericin B in both *S. cerevisiae* and *C. albicans* at concentrations far below their MIC_50_’s (Fig 6A and B). As the potentiation applies broadly to antifungals with varied mechanisms of action and to metals, we reason that this is consistent with a broad-based disruption of the ability of fungal cells to detoxify drugs. Our initial metabolic studies showing that CoA levels are increased (Fig. 6E) following treatment with PZ-2891 suggest that the compound’s activity is achieved not through direct inhibition of Cab1 activity but through inhibition of either Acs1 or Acs2. Meanwhile, since treatment with PZ-2891 does not inhibit growth in glucose-rich media, and *ACS2* knockouts or mutants are inviable in such conditions, this leaves Acs1 as the primary candidate for PZ-2891’s target in fungi. Our biochemical studies using purified yeast Acs1 demonstrated that PZ-2891, like AR-12, inhibits this enzyme’s activity (Fig. 6F). In clinical studies, an analog of PZ-2891, BBP-671 (NCT04836494), has been found to be largely safe with limited adverse events reported in humans ^36,37^. Thus, this class of small molecules may hold promise as the first antifungal adjuvants to enhance the potency of current drugs against drug-sensitive and -resistant strains while also lowering their toxicity.

In summary, our study shows that modulation of PanK activity results in impaired vacuolar homeostasis and xenobiotic detoxification, which in turns leads to enhanced fungal susceptibility to antifungal drugs. Therefore, compounds that target PanK or other key enzymes in the PCA pathway represent a promising path towards the development of novel therapeutic strategies to help combat the emerging threats posed by multi-drug resistant fungi and possibly other eukaryotic pathogens.

## Materials and Methods

### Yeast strains and vectors

Yeast strains used in this study are shown in Table 1.

**Table 1:**
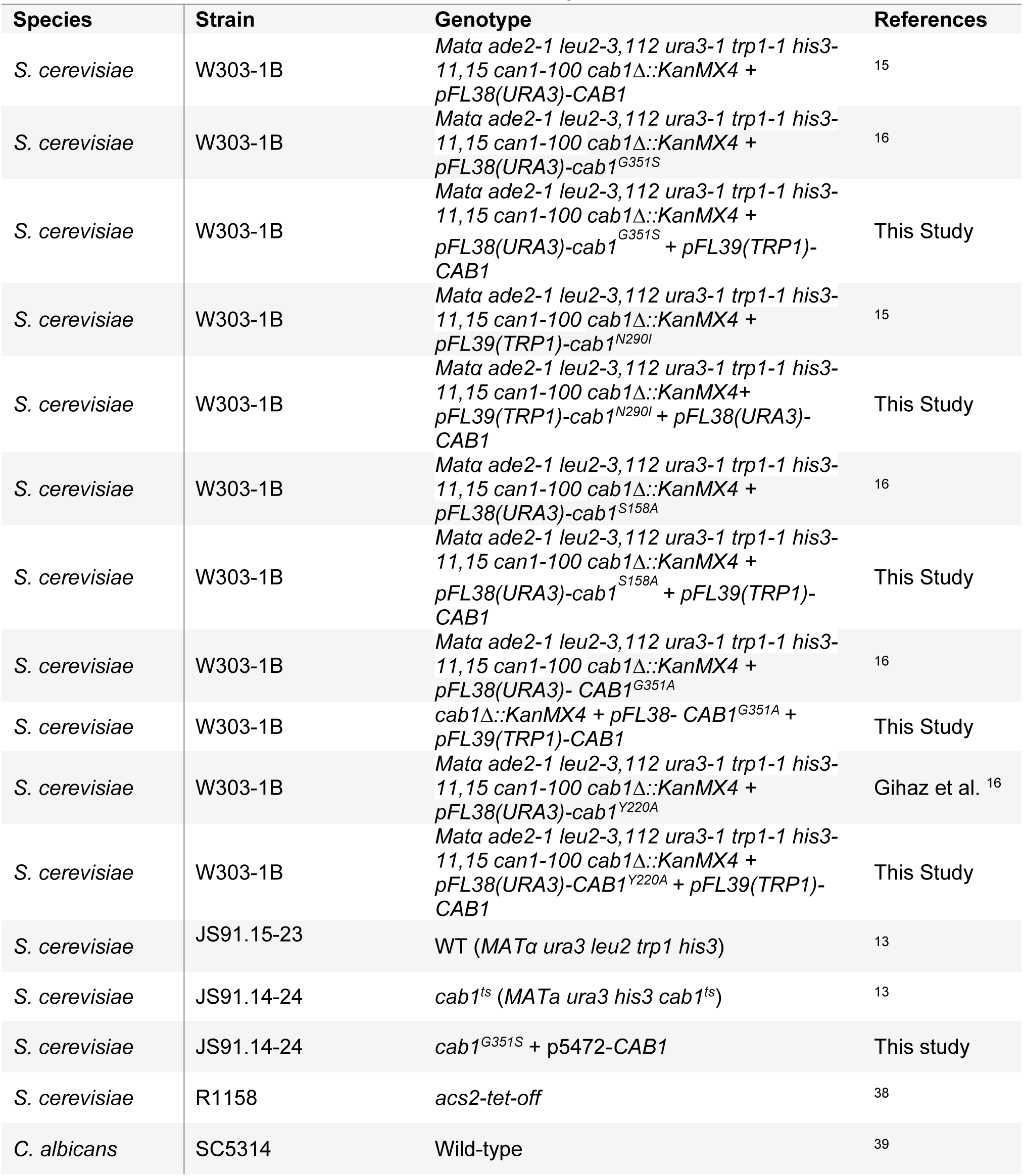
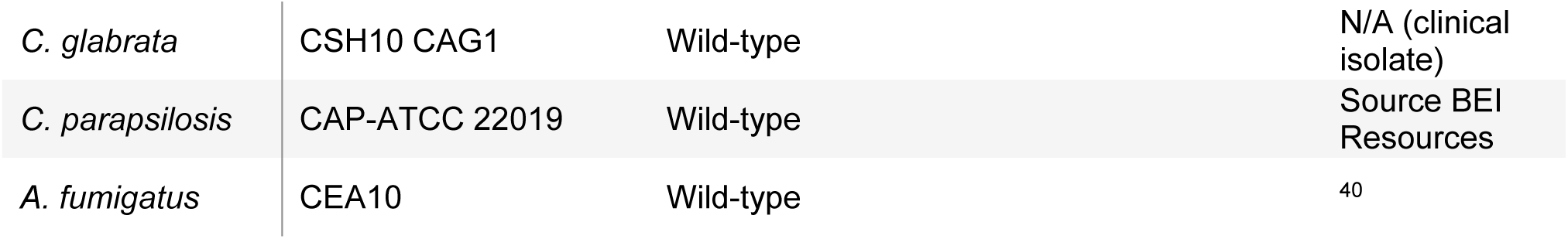
Yeast strain and their vectors used in this study.

### Selection of yeast strains carrying *cab1* mutations

cab1Δ/pFL38-*CAB1* wild type and mutant strains were generated using plasmid shuffling as previously described from the parent strain *cab1*Δ/pFL39-*cab1^G351S^* ^16^. Add-back strains were generated by introducing pFL39-*CAB1* vector into yeast recipient strains.

### Growth assays

Yeast strains (WT and *cab1* mutants) were grown overnight at 30°C in YPD medium and harvested (700 × g for 5 mins at 4°C), washed with water, and resuspended in 0.9% NaCl solution at OD_600_ of 0.5. Serial 10-fold dilutions were made and 5 µL of cell suspensions were spotted on YPD agar plates containing various antifungals (amphotericin B, caspofungin, fluconazole, terbinafine, hygromycin and cycloheximide). For the respiratory growth assay, YP medium supplemented with ethanol, lactic acid or glycerol were used. Plates were incubated at 30°C and the growth was monitored by image scan using the device ChemiDoc MP (Bio-Rad) every 24 hours. For the liquid growth assay, the yeast strains were pre-grown as above and then diluted into 3 mL of yeast rich media supplemented with either 2% glucose (YPD), 2% glycerol (YPG) or 2% lactate (YPL) liquid media at the concentrations of 10 cells per μL and incubated at 30°C by shaking at 230 rpm and the cell growth was monitored by optical density (OD_600_). *A. fumigatus* growth was examined on GMM (1% glucose, 6 g/L NaNO3, 0.52 g/L KCl, 0.52 g/L MgSO4•7H2O, 1.52 g/L KH2PO4 monobasic, 2.2 mg/L ZnSO4•7H2O, 1.1 mg/L H3BO3, 0.5 mg/L MnCl2•4H2O, 0.5 mg/L FeSO4•7H2O, 0.16 mg/L CoCl2•5H2O, 0.16 mg/L CuSO4•5H2O, 0.11 mg/L (NH4)6Mo7O24•4H2O, and 5 mg/L Na4EDTA; pH 6.5).

### Electron Microscopy Analysis EM

Yeast strains (WT and *cab1* mutants) were grown overnight at 30°C in YPD medium, harvested, and refreshed in YP media with 2% glycerol until reached OD_600_ of 1. The cells were harvested, washed, and used for high pressure freezing and freeze substitution for electron microscopy analysis. Unfixed samples were high pressure frozen using a Leica HMP100 at 2000 psi. The frozen samples were then freeze substituted using a Leica Freeze AFS unit starting at -95°C using 0.1% uranyl acetate in acetone for 50 h to -60°C, then rinsed in 100% acetone and infiltrated over 24 h to -45°C with Lowicryl HM20 resin (Electron Microscopy Science). Samples were placed in gelatin capsules and UV hardened at -45°C for 48 h. The blocks were allowed to cure for a further few days before trimmed and cut using a Leica UltraCut UC7. The 60nm sections were collected on formvar/carbon coated nickel grids and contrast stained using 2% uranyl acetate and lead citrate. The 60nm sections on grids were viewed FEI Tecnai Biotwin TEM at 80Kv. Images were taken using AMT NanoSprint15 MK2 sCMOS camera.

### Pantothenate utilization assay using ^14^C -labeled PA

Pantothenate utilization assay using labeled PA was performed as previously described ^13,14^. Briefly, cell-free extracts from yeast producing Cab1 variants were obtained by homogenization, followed by centrifugation at 700 x g for 5 min. The 40 μL enzyme reaction contained reaction buffer (100 mM Tris HCl, 2.5 mM MgCl_2_, 2.5 mM ATP, pH 7.4), D-[1-^14^C] pantothenate (2 nmol, 0.1 µCi), and 144 μg cell-free extracts. The lysates total protein content was determined using the Bradford assay. The reaction was done at 30 °C for 10 min following the addition of 4 µL of 10% acetic acid to stop the reaction. The reaction mixture was spotted on a DE-81 filter (0.6 mm in diameter) placed within a spin column with a 2 mL collection tube.

Following 5 min incubation, the spotted filters were centrifuged for 20 s at 700 x g, washed twice with 1% acetic acid in ethanol, and collected for liquid scintillation spectrometry.

### Cellular CoA determination

The determination of cellular CoA levels in yeast strains producing harboring different Cab1 variants was done using soluble metabolites extractions from *S. cerevisiae* as previously described ^41^. Metabolites extracts were then used in Coenzyme A detection kit (Sigma) to quantify cellular CoA.

### Cellular cysteine determination

The determination of cellular cysteine levels in yeast strain producing different Cab1 variants was done using soluble metabolites extractions from *S. cerevisiae* as previously described ^41^. Metabolites extracts were then used in fluorometric cysteine assay kit (Abcam).

### RNA sequencing and data analysis

RNA samples from yeast strain producing different Cab1 variants were extracted using YeaStar RNA kit (Zymo Research).

*RNA Seq Quality Control:* Total RNA quality is determined by estimating the A260/A280 and A260/A230 ratios by nanodrop. The RNA integrity is determined by resolving an aliquot of the extracted RNA on Agilent Bioanalyzer gel, which measures the ratio of the ribosomal peaks. Samples with RNA integrity number (RIN) values of 7 or greater are recommended for library preparation.

*RNA Seq Library Prep:* The mRNAs are purified from approximately 200ng of total RNA with oligo-dT beads and sheared by incubation at 94 °C in the presence of Mg (Kapa mRNA Hyper Prep). Following first-strand synthesis with random primers, second strand synthesis and A-tailing are performed with dUTP for generating strand-specific sequencing libraries. Adapter ligation with 3’ dTMP overhangs are ligated to library insert fragments. Library amplification amplifies fragments carrying the appropriate adapter sequences at both ends. Strands marked with dUTP are not amplified. Indexed libraries that meet appropriate cut-offs for both are quantified by qRT-PCR using a commercially available kit (KAPA Biosystems) and insert size distribution determined with the LabChip GX or Agilent Bioanalyzer. Samples with a yield of ≥0.5 ng/μL are used for sequencing. *Flow Cell Preparation and Sequencing:* Sample concentrations are normalized to 1.2 nM and loaded onto an Illumina NovaSeq flow cell at a concentration that yields 25 million passing filter clusters per sample. Samples are sequenced using 100bp paired-end sequencing on an Illumina NovaSeq according to Illumina protocols. The 10bp unique dual index is read during additional sequencing reads that automatically follow the completion of read 1. Data generated during sequencing runs are simultaneously transferred to the YCGA high-performance computing cluster. A positive control (prepared bacteriophage Phi X library) provided by Illumina is spiked into every lane at a concentration of 0.3% to monitor sequencing quality in real time.

*Data Analysis and Storage:* Signal intensities are converted to individual base calls during a run using the system’s Real Time Analysis (RTA) software. Base calls are transferred from the machine’s dedicated personal computer to the Yale High Performance Computing cluster via a 1 Gigabit network mount for downstream analysis. Primary analysis - sample de-multiplexing and alignment to the human genome - is performed using Illumina’s CASAVA 1.8.2 software suite. The data are returned to the user if the sample error rate is less than 2% and the distribution of reads per sample in a lane is within reasonable tolerance. Data is retained on the cluster for at least 6 months, after which it is transferred to a tape backup system.

*Data analysis:* Partek Flow was used to organize and process fastq files from paired-end sequencing. Paired-end reads were trimmed for quality (Q-Score > 20, Min read length 25) and adapters were removed using FastQC, then aligned to the *S. cerevisiae* W303-1B genome (Accession: JRIU00000000; ATCC Number: 200060, downloaded from SGD) using STAR v2.7.8a. Reads were normalized via trimmed mean of M values (TMM) ^42^. and further processed in Excel, where the three biological replicates per condition (WT, three mutants, and three mutants with addback) were grouped to generate means and standard deviations for all genes in each of the 7 conditions. Log fold changes in **Fig. 4E** correspond to the log base 2 of the ratio between one of 6 experimental conditions and the WT control condition. To quantify the degree of confidence of rescue of a gene differentially expressed in a mutant by its addback, the following decision matrix was utilized. First, the p-value between the wild type and mutant expression was computed using the standard deviations and means of the WT (N=3) and mutant (N=3 for each mutant) expression. Then, the p-value between wild type and complemented mutant was computed. “Large” circles in **Fig. 4E** corresponds to genes for which the first p-value<0.05, but the second p-value ≥0.05, indicating the null hypothesis of equal expression was rejected for the mutant, but not rejected for the addback at p=0.05. “Small” circles correspond to genes for which either of these criteria were not met.

### Seahorse analysis

The oxygen consumption rate (OCR) of yeast strains expressing different *CAB1* variants was determined using Seahorse 96X and Mito Stress kit. Yeast strains (WT and *cab1* mutants) were grown overnight at 30°C in YPD medium, harvested, and refreshed in SC medium supplemented with 2% glucose until reached OD_600_ of 0.6. Then, cells were harvested, washed and seeded (6 X 10^4^ cells per well) in Seahorse XFp plates coated with poly-Lysine (50 µL of 0.1 mg/mL). A minimum of 8 technical replicates were performed for each experiment at 30°C. The seeded plate was centrifuged at 500 rpm for 5 min to promote yeast adhesion and the plate was rested for 30 min at RT. A soaked and calibrated Seahorse XF96 Sensor Cartridge was prepared before loading into the Seahorse XF96 analyzer (Agilent) which determined the cells basal OCR and following the injection of mitochondrial uncoupling drugs; oligomycin (5 µM), carbonyl cyanide-4 (trifluoromethoxy) phenylhydrazone (FCCP) (10 µM), antimycin A (10 µM), and rotenone (5 µM). The readouts were normalized using nuclear Hoechst staining for the immobilized yeast cells.

### Yeast growth in the presence of antifungal drugs, inhibitors, and potentiators

To investigate the effect of common AFDs (amphotericin B, caspofungin, fluconazole, and terbinafine) in combination with compounds and potentiators (concanamycin A, doxycycline, PZ-2891, α-PanAm, AR-12) the yeast growth was monitored using liquid assay in a 96-well plate. Overnight yeast precultures (WT strains or acs-Tet-Off when mentioned) were prepared in YPD medium at 30°C. Cells were washed and refreshed in YPD until reaching OD_600_ of 0.6. In a 96-well plate, cells (10^3^ cells/mL, 100 µL final volume) were treated with decreasing concentrations (two-fold dilutions) of AFDs, and different dosages of compounds and potentiators. For reference, amorolfine (200 µM) and DMSO (0.6%) were used as positive and negative controls to determine 100% and 0% growth inhibition, respectively. Plates were incubated at 30°C. Optical density measurements were taken using a BioTek SynergyMx microplate reader every 12 h. Data are shown as mean ± SD of four independent experiments. Growth curves where visualized and determined from a sigmoidal dose-response curve using GraphPad Prism version 9.5.1 (GraphPad Software, San Diego, CA). Statistical significance was determined using t-test (*p=0.05*) with GraphPad Prism.

### Fluorescence microscopy of cell organelles

To visualize vacuolar structure, the yeast strains were grown to OD of 1 at 600 nm, harvested and resuspend cells at 10^6^ cells/mL in 10 mM HEPES buffer, pH 7.4, containing 5% glucose. CellTracker™ Blue CMAC was added to the cell suspension to a final concentration of 100 µM. The cells were incubated at room temperature for 30 minutes and the stained cells were visualized by fluorescence microscopy.

### Reactive Oxygen Species (ROS) content

Reactive oxygen species (ROS) in the *cab1* mutants was determined by change of oxidative status of fluorescence dye caused by ROS inside of the cell. ROS oxidize dihydrorhodamine 123 (DHR123; Sigma-Aldrich®, Darmstadt, Germany), which in turn produces green, fluorescent R123. To monitor ROS, cells were pre-grown overnight at 30 °C in YPD to the OD_600_ of 0.5 ∼1.0 and the cells were diluted to the OD of 0.4 and loaded with 1.25 µg/mL of DHR123 for 2 h at 30 °C. At the end of the incubation time, cells were harvested (2 min at 9,000 x g) and re-suspended in water at the OD_600_ of 0.05, and the fluorescence was quantified by a plate reader. For each sample, 100 µL of cell suspension was added into each well and the fluorescence was measured (excitation/emission spectra of 488/530 nm). Emission values from the control cells untreated with the dye were used as background for each strain. ROS generation for each *cab1* strains was measured as the percentage of fluorescence emission obtained from the *cab1Δ* strain harboring WT *CAB1* gene.

### PanK activity of recombinant Cab1 in the presence of PZ-2891 and AcCoA

His-tagged Cab1 recombinant enzyme was produced and purified as was previously described ^16^. A Kinase-Glo (Promega) assay kit for kinase activity was used to determine the activity of the purified PanK under different conditions ^16^.

### Vacuolar visualization and cell size determination

To determine the ratio of vacuolar area over cell area, different yeast strains were stained with CellTracker™ Blue CMAC as explained in the methods above. Images were captured using fluorescence microscope and analyzed using Image J software. The cell surface area (in square pixel) and vacuolar surface area (in square pixel) were calculated in Image J and percentage of vacuolar area/cell area was calculated. A total of 100 cells were analyzed from each yeast strain. The data was plotted and analyzed in GraphPad Prism version 9.5.1 (GraphPad Software, San Diego, CA). Statistical significance was determined using Welch’s t-test with GraphPad Prism.

### Radial growth assay and AFD sensitivity assays with *A. fumigatus*

The radial growth measurements of *A. fumigatus* were performed as previously described ^16,43^. Briefly, 2 μL of a 2.5 X 10^6^ mL-1 conidial suspension of wild-type CEA10 *A. fumigatus* was point inoculated onto the center of a solid GMM in the absence or presence of 50 µM PZ-2891, 20 µg/mL caspofungin, and their combination. Plates were incubated for 96 h at 35°C, with colony diameters measured and photographed taken each day.

### Acetyl CoA synthetase (ACS) activity assay

The ACS assay was performed by monitoring formation of the adenyl acetate, the intermediate of the enzyme reaction, utilizing *S. cerevisiae* acetyl Coa synthetase (Sigma, A1765), following established protocols with some modifications ^26,29^. In a 100 µL reaction volume, composed of 100 mM potassium phosphate at pH7.5, 5 mM MgCl_2_, 2 mM ATP, 50 mM potassium fluoride, 10 mM reduced glutathione, 0.35 mM CoA, 10 mM potassium acetate, 200 mM neutralized hydroxylamine adjusted to pH7.3, 0.005 units of the enzyme, and the inhibitors (in 1% DMSO), the components were combined. The mixture was then incubated for 30 minutes at 37 °C. Termination of the reaction was achieved by addition of 50 µL of a solution containing ferric chloride (12 M) and trichloroacetic acid (12%). The resultant product, acethydroxamic acid, was quantified using a BioTek SynergyMx microplate reader at OD_540_. The background correction was performed by utilizing a blank reaction comprising all the reaction components, which was subsequently terminated using acidified ferric chloride solution, without undergoing any incubation time.

## Acknowledgments

This research was supported by an award to C.B.M. by the Blavatnik Family Foundation. CBM research is also supported by funds from NIH and the Steven & Alexandra Cohen Foundation. We thank Dr. Paola Goffrini for providing the yeast strain (*S. cerevisiae W303-1B cab1Δ*/pFL39-*cab1*^G351S^), Dr. Hans-Hoachim Schuller for providing the *cab1^ts^* mutant and its parental strain, Dr. Frederick Roth for providing the yeast efflux-deficient mutant and its parental strains, and Dr. Joseph C. Gennaro for his assistance with the analysis of RNAseq data. This project has also been funded in whole or in part with Federal funds from the National Institute of Allergy and Infectious Diseases, National Institutes of Health, Department of Health and Human Services, under Contract No. HHSN272201700059C.

## Author Contributions

Conceptualization, J.Y.C., S.G., M.M., P.S., P.H., J.C.G., K.F., and C.B.M.; Methodology, J.Y.C., S.G., M.M., P.S., E.M.A., and J.C.G.; Formal Analysis, J.Y.C., S.G., M.M., P.S., P.V., E.M.A., and J.C.G.; Investigation, J.Y.C., S.G., M.M., P.S., P.H., E.M.A., J.C.G., K.F., and C.B.M.; Resources, J.Y.C., S.G., M.M., P.S., P.H., E.M.A., J.C.G., K.F., and C.B.M.; Writing – Original Draft, J.Y.C., S.G., P.S., P.V., P.H., J.C.G., K.F., and C.B.M.; Writing – Review & Editing, J.Y.C., S.G., P.S., P.V., P.H., J.C.G., K.F., and C.B.M.; Visualization, J.Y.C., S.G., M.M., P.S., E.M.A., J.C.G., K.F., and C.B.M.; Supervision, P.H., K.F., and C.B.M.; Project Administration, P.H., J.C.G., K.F., and C.B.M.; Funding Acquisition, K.F. and C.B.M.

## Supplementary figure legends

**Fig. S1. Cellular CoA (A) and cysteine (B) levels in *cab1Δ* strains harboring various *CAB1* mutations.** CoA and cysteine levels were measured using the metabolite extracts from the yeast strains grown in the presence of 1 µM PA.

**Fig. S2. RNA-Seq analysis for cysteine and sulfur homeostasis genes expressed in *cab1Δ* strains.** The results are based on normalized TMM compared with the expression profile of the WT parent strain. The gene list with annotations shown in **Table S1**.

**Fig. S3. RNA-Seq analysis for PCA pathway genes expressed in *cab1Δ* strains harboring various *CAB1* mutations.** Large circles in corresponds to genes for which the p-value (WT vs. mutant) <0.05, but the p-value (WT vs. addback) >0.05, indicating the null hypothesis of equal expression was rejected for the mutant, but not rejected for the addback at p=0.05. Small circles correspond to genes for which either of these criteria were not met. The full gene list is shown in **Table S1**.

**Fig. S4. Effect of modulation of downstream steps from the PCA pathway on the growth of *S. cerevisiae*.** *S. cerevisiae* (WT or acs2-tetoff mutant, as mentioned) cells were inoculated in the presence or absence of rising concentration of **A)** doxycycline, **B)** AR-12, and **C)** concanamycin A, at 30°C for 24-48 h. The growth was normalized to DMSO treated wells (no drug=100% growth) and 200 µM amorolfine well (0% growth).

**Fig. S5. PA utilization in the *cab1Δ* strains harboring various *CAB1* mutations.** Cell free extracts of yeast expressing cab1 mutants were used to measure the endogenous PA utilization of *cab1* variants using D-[1-^14^C] pantothenate as a substrate for 10 min at 30°C. The extracts PA utilization was measured in the absence or presence of 20 µM PZ-2891 (hPanK3 activator), α-PanAm (known Cab1 inhibitor), and YU385599 (reported Cab1 inhibitor).

**Fig. S6. Potentiation of PZ-2891 on antifungal susceptibility in different yeast species. A)** caspofungin efficacy in *S. cerevisiae* with PZ-2891. *S. cerevisiae* cells were inoculated in the presence or absence PZ-2891, in combination with caspofungin treatment at 30°C for 24-48 h. The growth was normalized to DMSO treated wells (no drug=100% growth) and 200 µM amorolfine well (0% growth). For these assays, t-test was done among the mentioned groups (*p=0.05*). **B)** Potentiation of terbinafine efficacy in *C. albicans* with PZ-2891. *C. albicans* spotting growth assays were performed when cells were inoculated into YPD overnight, harvested, washed, and re-suspended in 0.9% NaCl. Serial dilutions of cells were spotted onto YPD plates containing terbinafine (20 µg/mL) in the presence or absence of PZ-2891 (50 µM) at 30°C for 4 days. **C)** Average growth rate (based on colony diameter) of *A. fumigatus* colonies in the presence or absence of PZ-2891 (50 μM) in combination with caspofungin treatment (20 μg/ml). The results were calculated after 72 h of growth.

**Fig. S7. PZ-2891 does not have inhibitory effect on either *S. cerevisiae* Cab1 enzymatic activity or *S. cerevisiae* cell growth. A)** Dose-response curve for AcCoA effect on recombinant Cab1 enzyme activity. **B)** Dose-response curve for PZ-2891 effect on recombinant Cab1 enzyme activity**. C)** Dose-response curve for AcCoA effect on recombinant Cab1 enzyme activity in the absence of presence of PZ-2891 or YU385599 (inhibitor control)**. D)** Liquid growth assay for *cab1Δ* strains harboring various *CAB1* mutations in the presence or absence of 1-100 µM PZ-2891.

