## Supplementary figures and images for "Vitamin B5 Metabolism is Essential for Vacuolar and Mitochondrial Integrity and Xenobiotic Detoxification in Fungi"

### Supplemental Figures

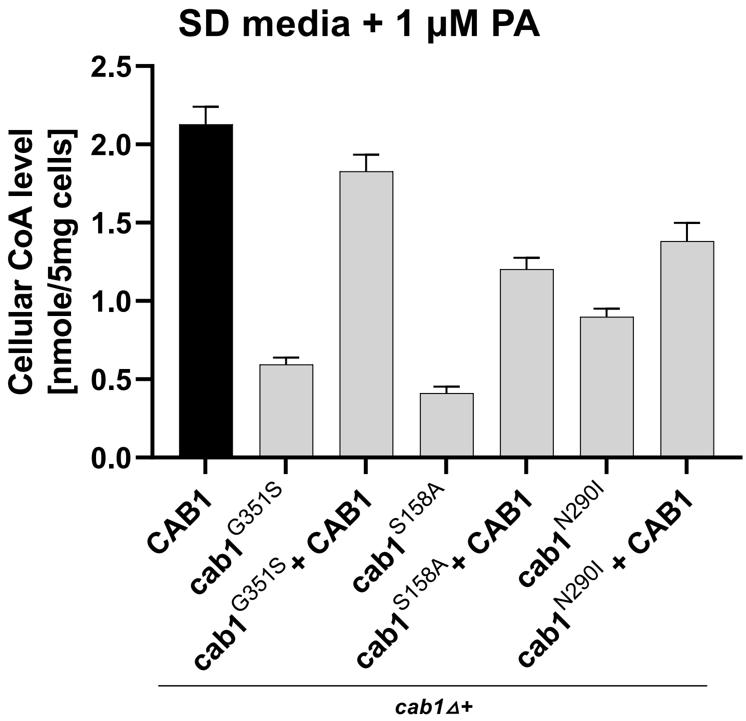

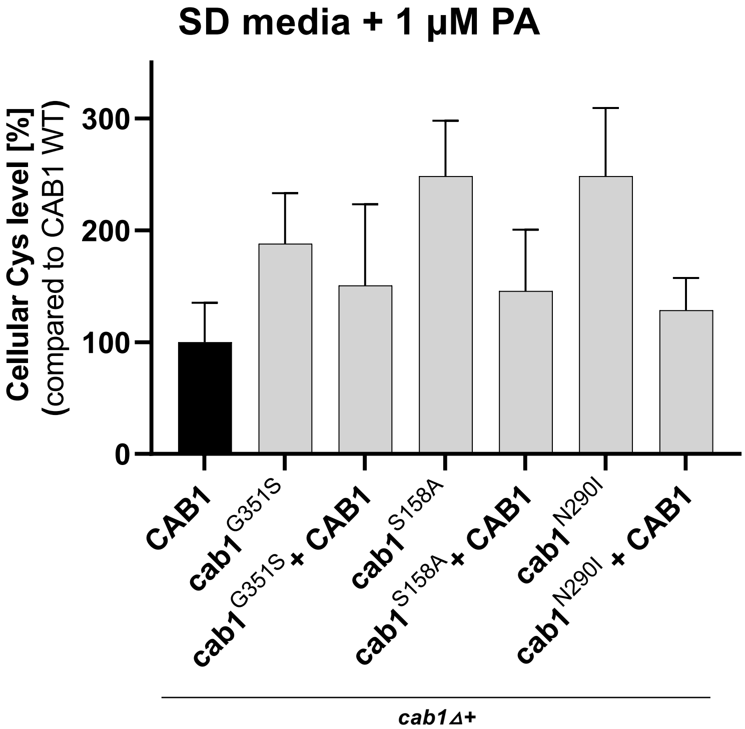

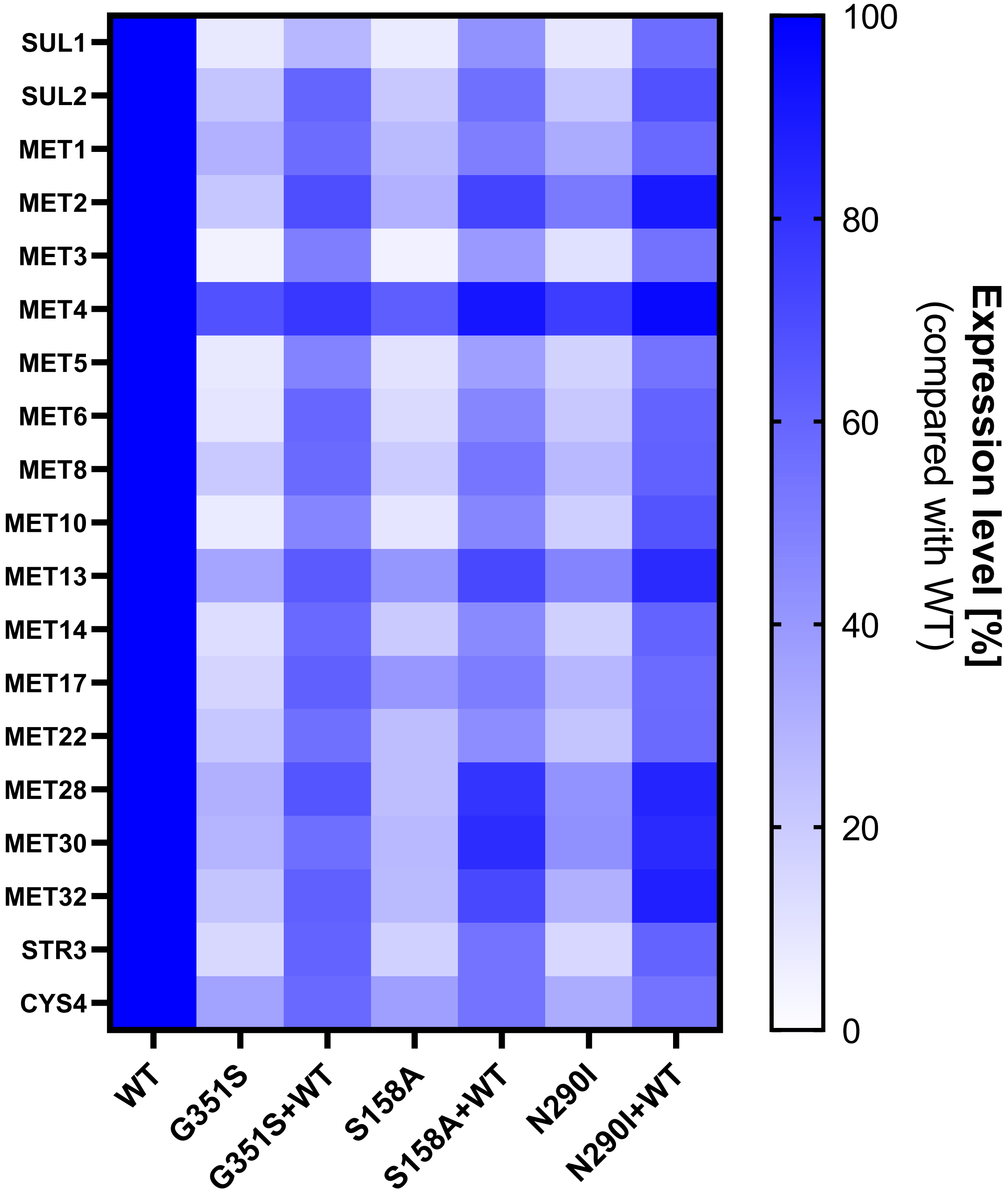

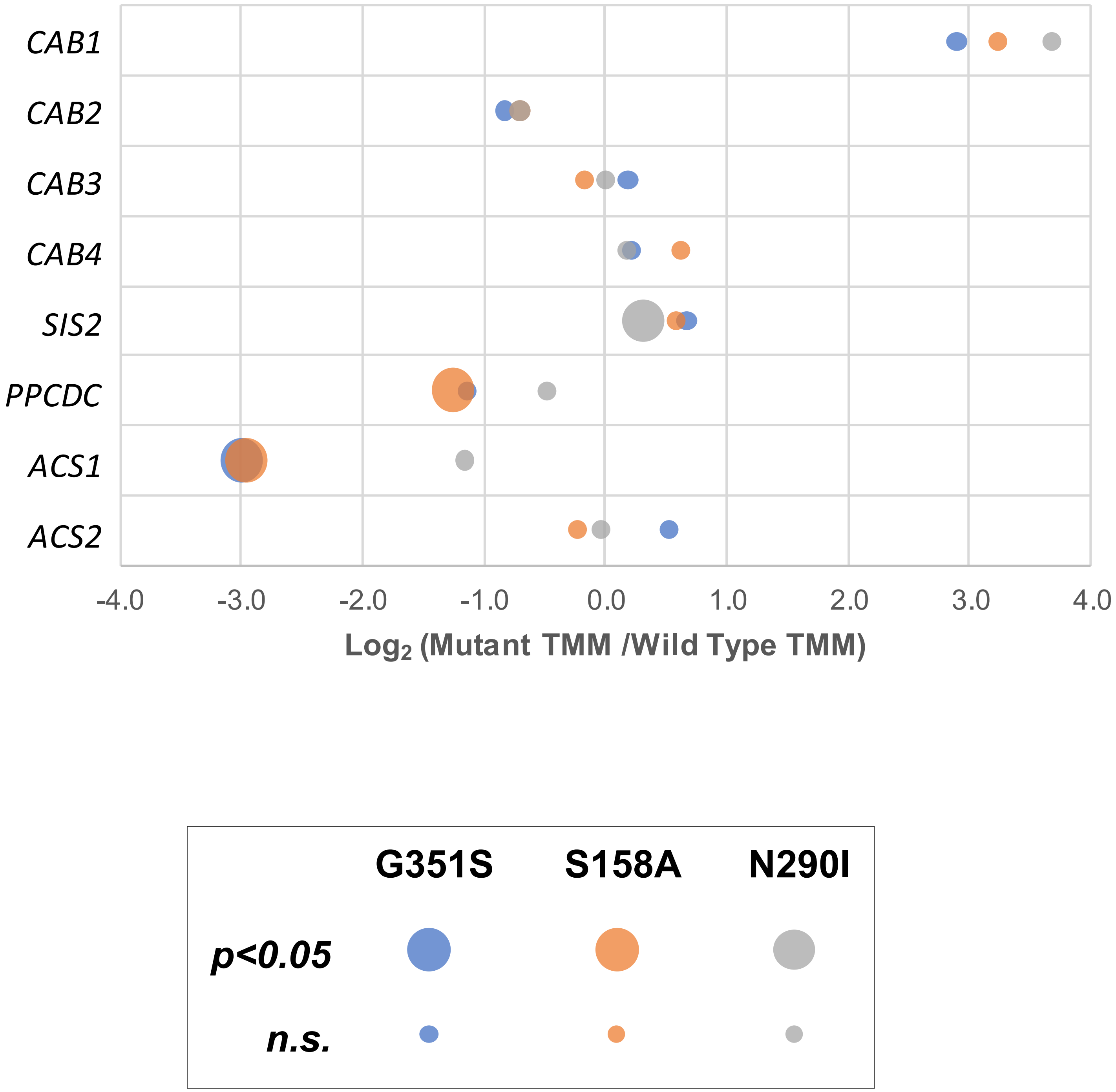

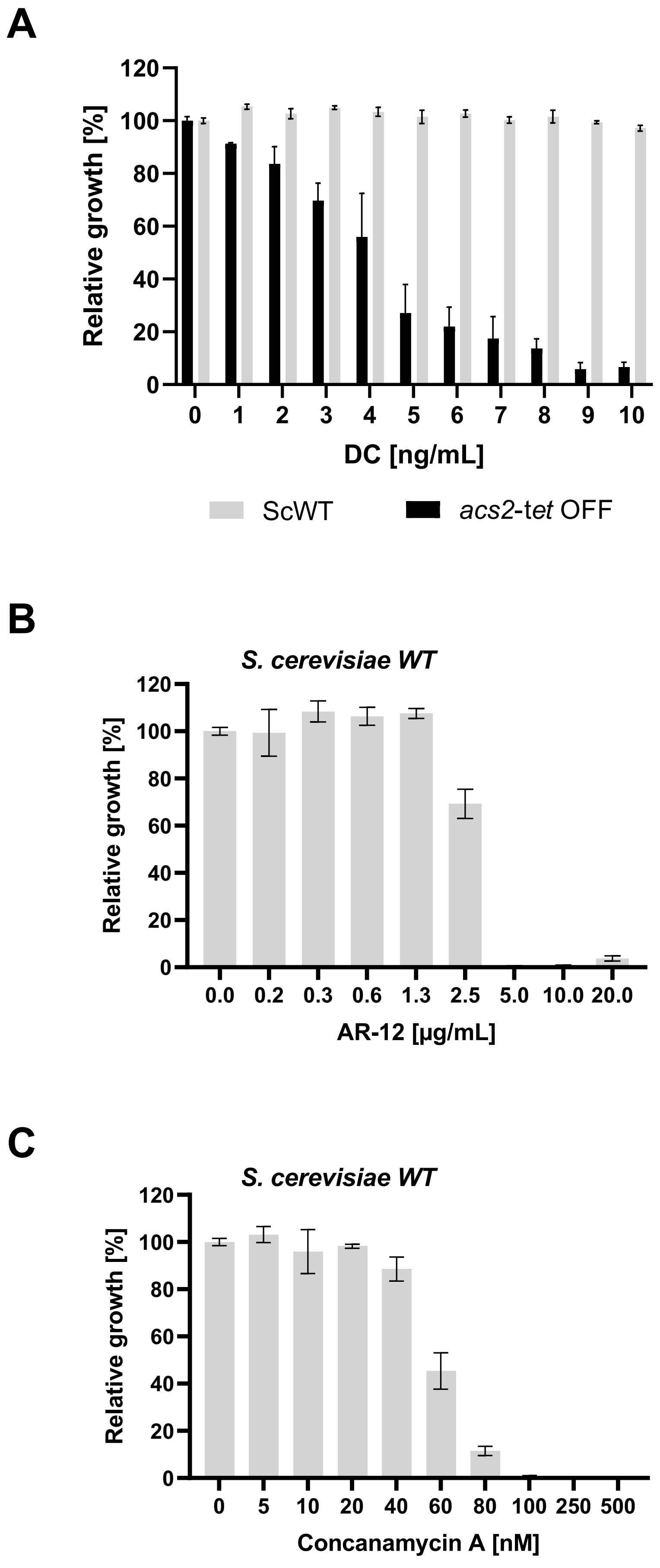

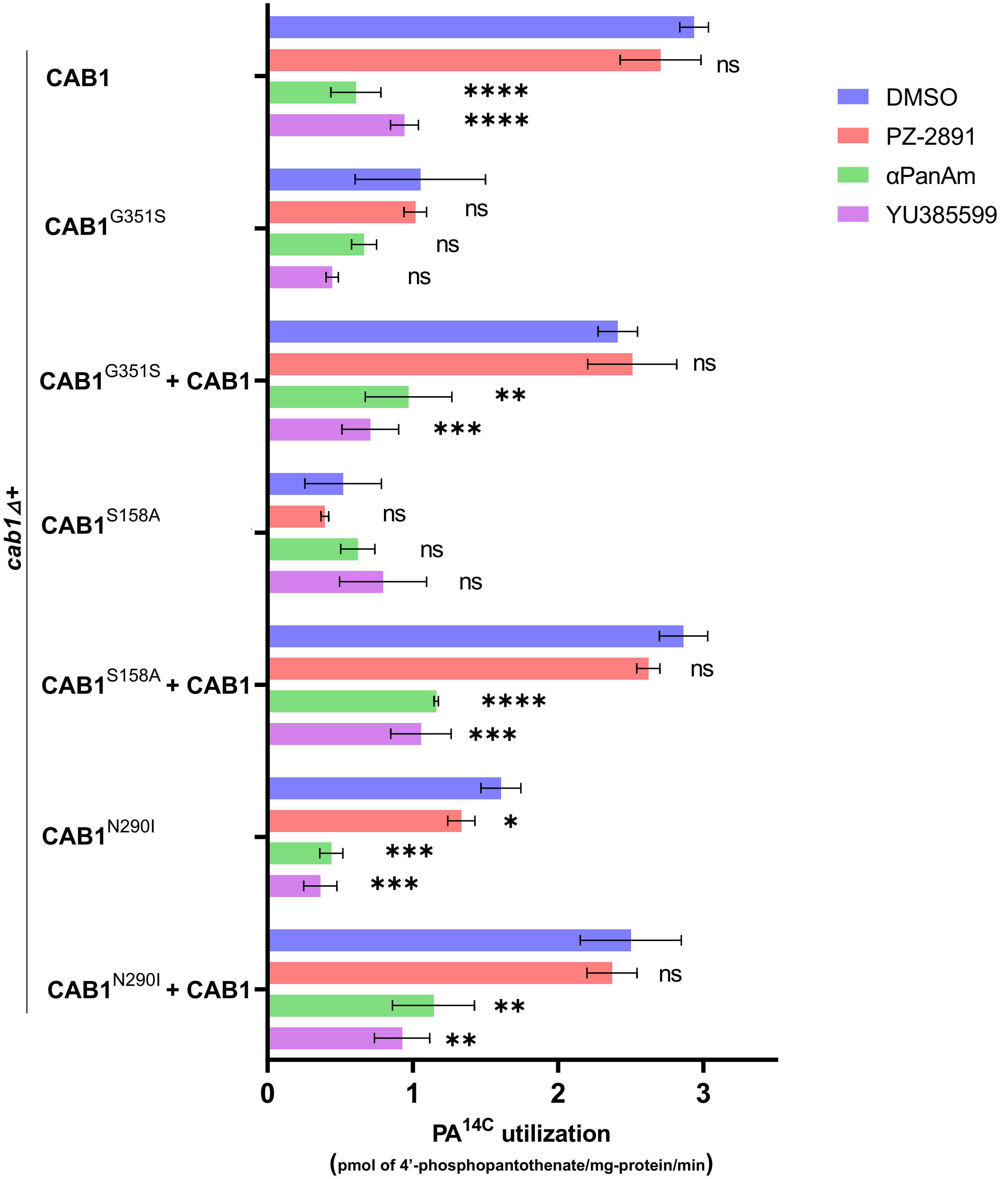

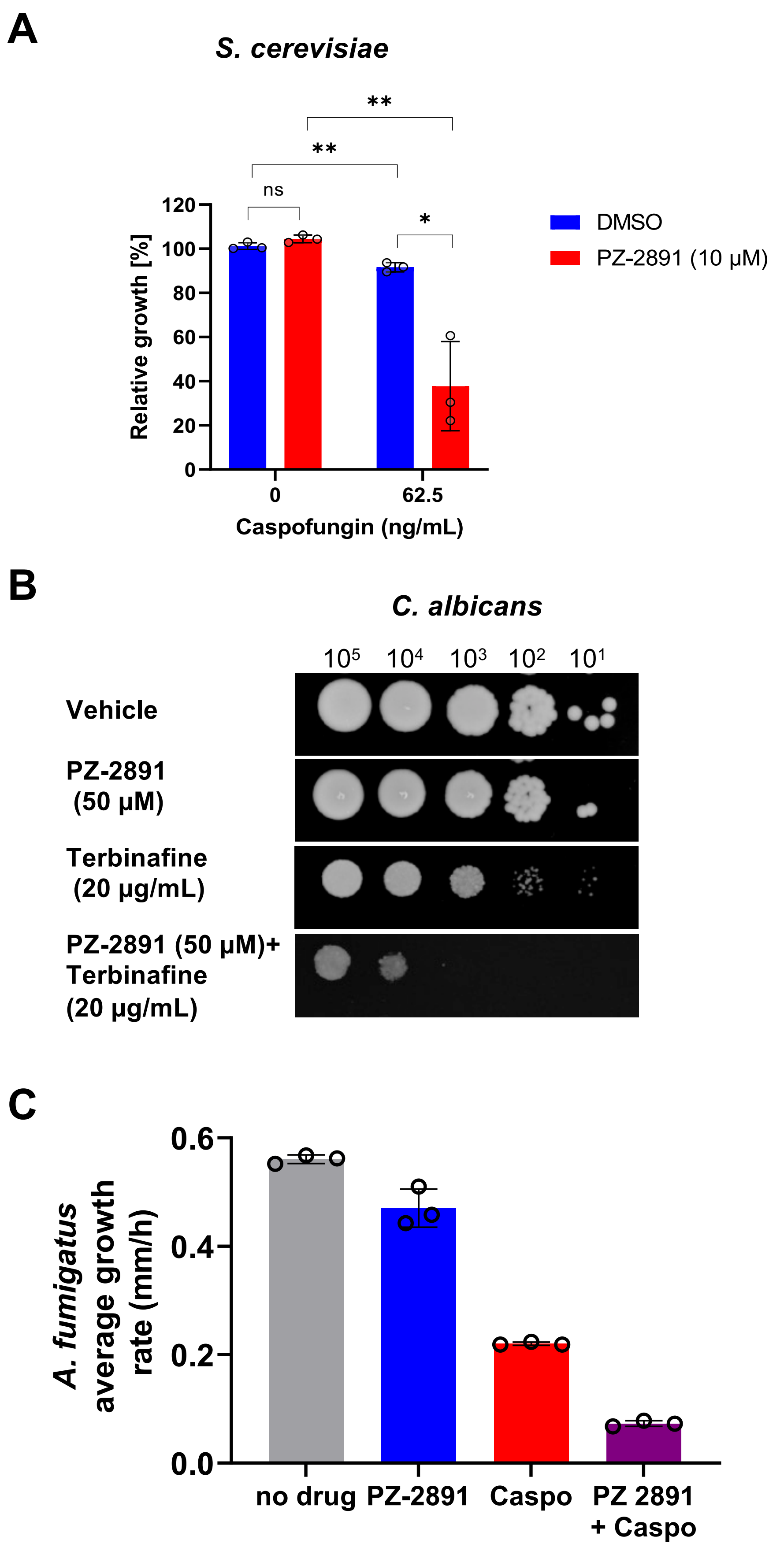

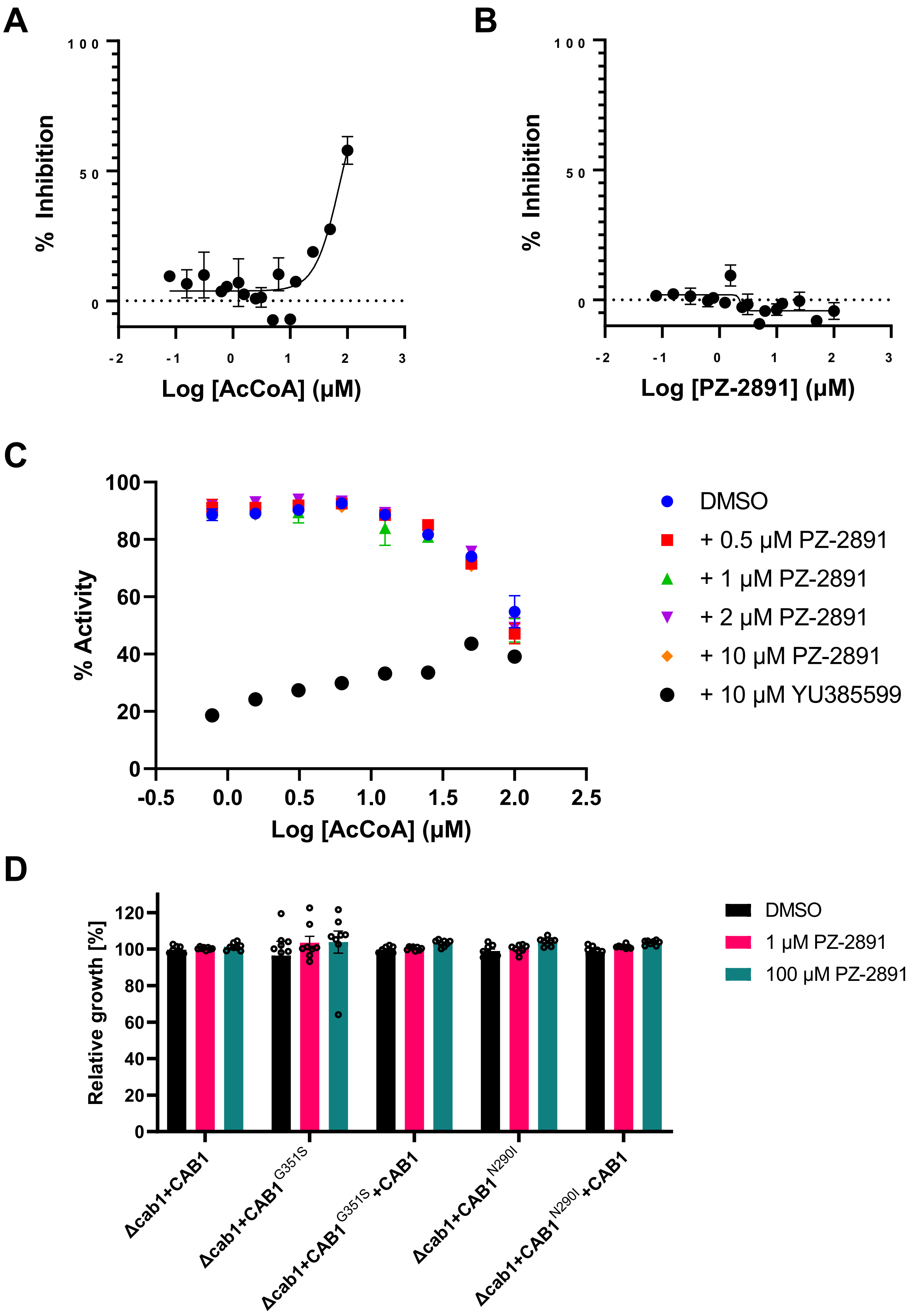
