## Supplemental Table 1 for "Vitamin B5 Metabolism is Essential for Vacuolar and Mitochondrial Integrity and Xenobiotic Detoxification in Fungi"

### Table S1- Down regulated genes list associated with *CAB1* deficiency

| Systematic name | Standard name and aliases | Short description |
|---|---|---|
| YDR531W | CAB1,pantothenate kinase | Pantothenate kinase, ATP:D-pantothenate 4'-phosphotransferase |
| YIL083C | CAB2,phosphopantothenate--cysteine ligase CAB2 | Phosphopantothenoylcysteine synthetase (PPCS) |
| YKL088W | CAB3,phosphopantothenoylcysteine decarboxylase complex subunit CAB3 | Subunit of PPCDC and CoA-SPC complexes involved in CoA biosynthesis |
| YGR277C | CAB4,putative pantetheine-phosphate adenylyltransferase | Subunit of the CoA-Synthesizing Protein Complex (CoA-SPC) |
| YKR072C | SIS2,HAL3,phosphopantothenoylcysteine decarboxylase complex subunit SIS2 | Negative regulatory subunit of protein phosphatase 1 (Ppz1p) |
| YOR054C | VHS3,YOR29-05,phosphopantothenoylcysteine decarboxylase complex subunit VHS3 | Negative regulatory subunit of protein phosphatase 1 Ppz1p |
| YAL054C | ACS1,FUN44,acetate--CoA ligase 1 | Acetyl-coA synthetase isoform |
| YLR153C | ACS2,acetate--CoA ligase ACS2 | Acetyl-coA synthetase isoform |
| YBR294W | SUL1,SFP2,sulfate permease | High affinity sulfate permease of the SulP anion transporter family |
| YLR092W | SUL2,sulfate permease | High affinity sulfate permease |
| YKR069W | MET1,MET20,uroporphyrinogen-III C-methyltransferase | S-adenosyl-L-methionine uroporphyrinogen III transmethylase |
| YNL277W | MET2,homoserine O-acetyltransferase | L-homoserine-O-acetyltransferase |
| YJR010W | MET3,sulfate adenylyltransferase | ATP sulfurylase |
| YJR137C | MET5,ECM17,sulfite reductase (NADPH) subunit beta | Sulfite reductase beta subunit |
| YER091C | MET6,5-methyltetrahydropteroyltriglutamate-homocysteine S-methyltransferase | Cobalamin-independent methionine synthase |
| YBR213W | MET8,bifunctional precorrin-2 dehydrogenase/sirohydrochlorin ferrochelatase MET8 | Bifunctional dehydrogenase and ferrochelatase |
| YFR030W | MET10,sulfite reductase subunit alpha | Subunit alpha of assimilatory sulfite reductase |
| YGL125W | MET13,MET11,MRPL45,methylenetetrahydrofolate reductase (NAD(P)H) MET13 | Major isozyme of methylenetetrahydrofolate reductase |
| YKL001C | MET14,adenylyl-sulfate kinase | Adenylylsulfate kinase |
| YLR303W | MET17,MET15,MET25,bifunctional cysteine synthase/O-acetylhomoserine aminocarboxypropyltransferase MET17 | O-acetyl homoserine-O-acetyl serine sulfhydrylase |
| YOL064C | MET22,3'(2'),5'-bisphosphate nucleotidase,HAL2 | Bisphosphate-3'-nucleotidase |
| YIR017C | MET28 | bZIP transcriptional activator in the Cbf1p-Met4p-Met28p complex |
| YDR253C | MET32 | Zinc-finger DNA-binding transcription factor |
| YGL184C | STR3,cystathionine beta-lyase STR3 | Peroxisomal cystathionine beta-lyase |
| YGR155W | CYS4,NHS5,STR4,VMA41,cystathionine beta-synthase CYS4 | Cystathionine beta-synthase |
